# Root growth responses to mechanical impedance are regulated by a network of ROS, ethylene and auxin signalling in Arabidopsis

**DOI:** 10.1101/2020.09.01.277707

**Authors:** Amy G.R. Jacobsen, Jian Xu, Jennifer F. Topping, Keith Lindsey

## Abstract

- The growth and development of root systems, essential for plant performance, is influenced by mechanical properties of the substrate in which the plants grow. Mechanical impedance, such as by compacted soil, can reduce root elongation and limit crop productivity.
- To understand better the mechanisms involved in plant root responses to mechanical impedance stress, we investigated changes in the root transcriptome and hormone signalling responses of Arabidopsis to artificial root barrier systems *in vitro*.
- We demonstrate that upon encountering a barrier, reduced Arabidopsis root growth and the characteristic ‘step-like’ growth pattern is due to a reduction in cell elongation associated with changes in signalling gene expression. Data from RNA-sequencing combined with reporter line and mutant studies identified essential roles for reactive oxygen species, ethylene and auxin signalling during the barrier response.
- We propose a model in which early responses to mechanical impedance include reactive oxygen signalling that is followed by integrated auxin and ethylene responses to mediate root growth changes. Inhibition of ethylene responses allows improved growth in response to root impedance, a result that may inform future crop breeding programmes.

## Introduction

When growing through soils, plant roots must be able to respond to a range of environmental cues and rely on flexible growth to adapt to changing conditions. Establishment of robust root growth and adaptive root architecture is important for maintaining crop yields, and hence root traits are of interest to plant breeders (Gewin, 2010). Various stress conditions in soils limit root elongation, including insufficient nutrients, oxygen or water, and physical barriers (Bengough *et al*., 2006). As plant roots navigate the soil environment, they encounter physical barriers and must be able to adapt their growth in order to respond mechanical impedance. An increase in the mechanical strength of soils can occur as a result of drying and drought conditions, as there is a strong interaction between soil strength and water content (Whalley *et al*., 2005; Jin *et al*., 2013). The effect of increasing mechanical strength can be further exacerbated by soil compaction caused through the use of heavy farming machinery (Jin *et al*., 2013). Penetrometer resistance is commonly used as a measure of soil strength, and increased resistance correlates with reduced root elongation (Whitmore & Whalley, 2009; Bengough *et al*., 2011).

Mechanical impedance of the root also affects aerial growth. When roots are mechanically impeded shoot growth slows (Iijima & Kono, 1991; Roberts *et al*., 2002; Kobaissi *et al*., 2013; Potocka & Szymanowska-Pulka, 2018), and changes to leaf growth and morphology are also reported. Leaf number (Grzesiak, 2009), area (Alexander & Miller, 1991; Bingham *et al*., 2010; Kobaissi *et al*., 2013) and elongation rates (Young *et al*., 1997) decrease and stomatal closure has been observed (Roberts *et al*., 2002). There is therefore an agronomic relevance to understanding the response of roots to mechanical impedance, as the effects of soil drying and compaction can reduce crop yields (Whalley *et al*., 2008).

A diverse range of methods have been used to investigate the effect physically constraining root growth has on the morphology and architecture of root systems (Potocka & Szymanowska-Pulka, 2018), and many studies have focused predominantly on the morphological effect of mechanical impedance and barriers to root growth in crop species. It has previously been demonstrated that an increase in soil strength results in a decrease in root elongation in cotton (Taylor & Ratliff, 1969), maize (Bengough & Mullins, 1991), pea (Croser *et al*., 1999; Iijima & Kato, 2007) and tobacco (Alameda *et al*., 2012). In addition studies have demonstrated that in response to mechanical impedance, cell length is reduced and the length of the elongation zone shortened (Veen, 1982; Croser *et al*., 1999; Hanbury & Atwell, 2005; Okamoto *et al*., 2008). The slowed rate of elongation that occurs in mechanically impeded roots is therefore likely to be due to a reduced rate of cell elongation, as axial tension is increased by stiffening of the cell walls to reduce elongation (Bengough *et al*., 2006).

More recently, studies have begun focusing on the response of *Arabidopsis thaliana* in order to investigate in more detail the molecular basis of the root response to mechanical stimulus (Massa & Gilroy, 2003; Okamoto *et al*., 2008; Monshausen *et al*., 2009; Lee *et al*., 2020). Previous studies have sought to investigate the role of plant hormones and signalling pathways in roots in response to touch stimuli or mechanical impedance, however the exact nature of the signalling mechanism remains unknown. Evidence exists that both ethylene and auxin are likely to be involved in mediating the root response to mechanical impedance (Masle, 2002; Braam, 2005; Okamoto *et al*., 2008; Yamamoto *et al*., 2008; Lee *et al*., 2020). Changes in root morphology in mechanically impeded roots often resembles changes observed when roots are exposed to ethylene, an inhibition of root elongation and increase in the proliferation of root hairs (Masle, 2002; Buer *et al*., 2003). Classic studies have examined the response of roots to inclined, hard agar plates (1.5% as opposed to 1%) in order to examine thigmotropic responses in roots. Roots exhibit a wavy growth pattern due to the mechanical stimulus avoidance response (Okada & Shimura, 1990), and Buer *et al*. (2003) demonstrated that ethylene modulates this response. When Arabidopsis is grown in medium consisting of a normal layer and a harder layer, the root can show a bending response at the lower, harder layer. The bending or non-bending response of roots has been shown to depend on ethylene (Yamamoto *et al*., 2008). The role of ethylene signalling has also been demonstrated in continuously mechanically impeded *Arabidopsis* roots (Okamoto *et al*., 2008; Okamoto & Takahashi, 2019). It is possible that ethylene signalling mediates the response of roots to mechanical stress through co-action with auxin. It has previously been demonstrated that the effect of ethylene on root growth is mediated through regulation auxin biosynthesis and localisation of auxin transporters (Růžička *et al*., 2007; Strader *et al*., 2010).

Previous work also demonstrated that roots respond to obstacles through a combination of thigmotropic and gravitropic reactions (Massa & Gilroy, 2003). Auxin signalling has long been demonstrated to be involved in the gravitropic response of roots (Ottenschläger *et al*., 2003; Swarup *et al*., 2005; Muday & Rahman, 2008). It has also been proposed that the dynamic trafficking system of auxin and the auxin transport system in a key element in controlling tropic growth (Friml *et al*., 2002; Blilou *et al*., 2005; Pernisova *et al*., 2016). Three previous studies to study mechanical root impedance effects in Arabidopsis used either continuous mechanical impedance (horizontal root growth across the surface of a dialysis membrane; Okamoto *et al*., 2008) or the impedance of vertical root growth by a glass or metal barriers (Massa & Gilroy, 2003; Lee *et al*., 2020). Massa & Gilroy (2003) implicated barrier sensing via the root cap; Okamoto *et al*. (2008) demonstrated a role for ethylene signalling in inducing growth inhibition and radial thickening of roots; and Lee *et al*. (2020) showed a role for auxin transport via PIN2 to mediate root bending.

To understand better the network of signalling pathways involved the Arabidopsis root response to impedance, we used transcriptional profiling followed by experimental validation of gene expression data, with a focus on signalling pathway genes. We demonstrate that the early response involves an activation of reactive oxygen species (ROS) associated with ethylene and auxin signalling, and show that each is required for the impedance response.

## Materials and Methods

### Plant Material

Wildtype (WT) *Arabidopsis thaliana* seeds (Columbia (Col-0) ecotype), the auxin signalling mutant *axr1*, transport mutants *aux1-7* and *eir1-4*, and ethylene-insensitive *etr1* and *ein2* were from laboratory stocks. *atrbohF, atrobhD* and *atrbohD/F* mutants were from Prof. Alistair Hetherington (University of Bristol, UK). proCYCB1;2::CYCB1:2:GUS (Schnittger *et. al*, 2002) was from lab stocks. HyPer (Belousov *et al*., 2006) was from Prof. Marc Knight (Durham University, UK). DR5rev::3xVENUS-N7 (Brunoud *et al*., 2012) and R2D2 reporter lines (Liao *et al*., 2015) were from the Nottingham Arabidopsis Stock Centre (http://arabidopsis.info/). Seeds were stratified, surface sterilized and seedlings grown on sterile half strength Murashige and Skoog medium (1/2MS10) with 0.5% (w/v) agar, or 0.3% (w/v) or 1.2% (w/v) Phytagel depending on the assay, as described (Casson *et al*., 2002).

### Barrier Assays

We used three different barrier systems: a high density Phytagel (split medium) system, the use of plastic barriers and growth on the surface of a dialysis membrane, as follows.

#### 1. Split medium assay

Seedlings were grown in Magenta vessels (77 mm × 77 mm × 97 mm) containing a lower layer of 1.2% Phytagel 1/2MS10 medium, and once set, 0.3% Phytagel medium was poured on top. Seeds were placed just below the surface of the medium, to grow through the upper layer before reaching the barrier layer (Fig. S1a).

#### 2. Plastic and dialysis membrane barrier assays

One horizontal barrier impedance system was adapted from Massa & Gilroy (2003). Seeds were grown on vertical plates containing 1/2MS10 with 0.5% Phytagel. 1 x 2 cm plastic barriers of clear acetate were sterilised in 70% ethanol and at 6 days after germination (DAG) were placed horizontally directly in front of growing root tips (Fig. S1b); control roots had no barriers. For chemical treatments, seedlings were first germinated on standard 1/2MS10 medium and transferred at 6 DAG to the treatment plates before barrier placement. For some experiments, roots were grown along the surface of a dialysis membrane on the surface of a horizontal agar plate, to provide continuous impedance, as described (Okamoto *et al*. 2008; Fig. S1c).

For root architecture analysis, seedlings were photographed using a Zeiss STEMI SV8 dissecting stereomicroscope (Carl Zeiss, Cambridge, UK). Each growth assay was repeated 3 times with 15 individuals per treatment. All image analysis was carried out using ImageJ, and lateral root analysis used the Smart Root Plugin (Lobet *et al*., 2011). Root tip angle (RTA) was measured as the angle between the outer root tip and the horizontal barrier (Massa & Gilroy, 2003; Fig. S1d). For root hair length analysis, up to 20 mature root hairs were measured. The ImageJ straight-line tool was used to draw and measure a line from the quiescent centre (QC) to the proximal end of the meristem (defined as the first cell that was twice the length of the immediately preceding cell; González-García *et al*., 2011). Elongation zone size was measured from the end of the meristem to the first indication of emerging root hairs. Time-lapse imaging of roots used a Dino-lite microscope over 12 h, with images captured every 15 min. RTA and root growth was measured in each image from the point at which the root tip made contact with the barrier.

### Histochemistry

Histochemical staining of proCYCB1;2::CYCB1:2:GUS activity with clearing in chloral hydrate solution was as described (Topping & Lindsey, 1997). Number of dividing (GUS-positive) cells was calculated using the ImageJ ‘Cell Counter’ plugin. CellROX Deep Red dye (ThermoFisher, UK) revealed intracellular accumulation of ROS, and has previously been used for plant roots (Kováčik *et al*., 2014; Wright *et al*., 2016). Whole seedlings were stained with 1 μM dye for 30 min then washed twice with sterile distilled H_2_O.

### Confocal Microscopy

Roots were imaged using a Leica SP5 laser scanning confocal microscope (www.leica-microsystems.com) or Zeiss LSCM 880 (https://www.zeiss.com/-microscopy/int/home.html). Seedlings were fixed in 4% paraformaldehyde (PFA) before clearing with ClearSee (Kurihara *et al*., 2015) and staining with 0.1% (w/v) Calcofluor White for 30 min or unfixed tissues stained with 0.5% (w/v) propidium iodide (PI) solution for 90 s. Images were opened in LAS AF Lite software (v2.63 build 8173) or Zen Lite software. Images were taken from at least six individual roots for each analysis in Image J.

### Analysis of reporter fluorescence

The ImageJ polygon tool was used to delineate regions of interest (ROI) in the root tip. Fluorescence was measured either as mean green channel intensity or mean grey value. Mean green channel intensity was calculated in RGB images using the colour histogram tool. For grey values, all channels were exported as separate TIFFS and converted to 32-bit images and mean grey value measured using the ‘set measurement’ function in ImageJ. ROIs were selected using the ROI analyser tool and an RGB reference image with all channels, subtracting background from the value. For R2D2, reduction of the Venus signal relative to the tdTomato signal is a proxy for auxin accumulation (Liao *et al*., 2015). Separate TIFFS of the Venus and tdTomato channels were exported and converted to 32-bit in ImageJ. The Image Calculator function of ImageJ determined the ratio of Venus to tdTomato signals (https://imagej.nih.gov/ij/docs/menus/process.html#calculator). For HyPer analysis (Belousov *et al*., 2006), images were acquired at 488 nm (green channel) for analysis using Image Calculator. Images of CellROX DeepRed stained roots were obtained using both confocal and brightfield, using the electronically switchable illumination and detection module (ESID). Fluorescence was measured as mean grey value in 32-bit images containing only the DeepRed channel.

### RNA-sequencing

20 mg of tissue was ground in liquid nitrogen using TissueLyser II (QIAGEN, Manchester, UK) and RNA extracted using the Qiagen ReliaPrepTM RNA Tissue Miniprep System. RNA quality was determined using the NanoDrop ND-1000 spectrophotometer (ThermoFisher Scientific) and Agilent 2200 TapeStation. Libraries were constructed from 100 ng and 1 μg total RNA using the NEBNext UltraTM Directional RNA Library Prep Kit for Illumina for use with the NEBNext Poly(A) mRNA Magnetic Isolation Module (NEB, Hitchin, UK). mRNA was isolated, fragmented and primed, cDNA was synthesised and end prep was performed. NEBNext Adaptor was ligated and the ligation reaction was purified using AMPure XP Beads. PCR enrichment of adaptor ligated DNA was conducted using NEBNext Multiplex Oligos for Illumina (Set 1, NEB#E7335). The PCR reaction was purified using Agencourt AMPure XP Beads. Library quality was then assessed using a DNA analysis ScreenTape on the Agilent Technologies 2200 TapeStation. qPCR was used for sample quantification using NEBNext^®^ Library Quant Kit Quick Protocol Quant kit for Illumina. Samples were diluted to 10 nM. 7 μl of each 10 nM sample was pooled together and all were run on two lanes using an Illumina HiSeq2500 (DBS Genomics facility, Durham University). Approximately 30M unique paired-end 125bp reads were carried per sample. Primers were designed using Primer-BLAST (http://www.ncbi.nlm.nih.gov/tools/primer-blast/) and synthesised by MWG Eurofins (http://www.eurofinsdna.com/).

FastQC (https://www.bioinformatics.babraham.ac.uk/projects/fastqc/) was used to assess read quality and Trimmomatic (Bolger *et al*., 2014) was used to cut down and remove low quality reads. Salmon (Patro *et al*., 2017) was used for quasi-mapping of reads against the AtRTD2-QUASI (Brown *et al*., 2017; Zhang *et al*., 2017) transcriptome and to estimate transcript-level abundances. The tximport R package (Soneson *et al*., 2016) was used to import transcript-level abundance, estimate counts and transcript lengths, and summarise into matrices for downstream analysis in R. Before differential expression analysis, low quality reads were filtered out of the data set. Only genes with a count per million of 0.744 in 6 or more samples were retained. The DESeq2 (Love *et al*., 2014) R package was used to estimate variance-mean dependence in count data and test for differential expression (using the negative binomial distribution model). A padj-value of ≤0.05 and a log^2^fold change of ≥0.5 were selected to identify differentially expressed genes (DEGs). The 3D RNA-Seq online App (Guo *et al*., 2019; Calixto *et al*., 2018) was used for independent verification of estimated DEGs and for differential alternative splicing analysis. RNA-seq data are deposited in the Dryad Digital Repository.

## Results

### The root response to mechanical impedance involves radially asymmetric changes in cell expansion

In preliminary experiments to investigate the response of roots to a mechanical barrier, a two-layer gel system was used. Vertical root growth was established in a layer of soft medium (0.3% Phytagel) before hitting a harder lower layer (1.2% Phytagel), which creates mechanical impedance. Consistent with previous studies, primary root length significantly decreased on encountering the barrier, with median root length reducing from 15.8 mm (IQR = 4.0) to 10.6 mm (IQR = 3.4) (Fig. 1a). Root length between 4-7 days after germination (DAG) was significantly reduced on impedance, with a significant reduction occurring from 5 DAG (8.21 ± 0.24 mm compared to 5.65 ± 0.36 mm; Fig. 1g). The distance of root hair emergence from the root tip was also significantly reduced on encountering a barrier (Fig. 1b; Okamoto *et al*., 2008). The meristem length of impeded roots (7 DAG) was not significantly different to the control (Fig. 1c,e), but the elongation zone length was significantly reduced, from a median of 467 μm (IQR = 132) in the control to 362 μm (IQR = 123) in impeded roots (Fig. 1d,f).

**Figure 1.**
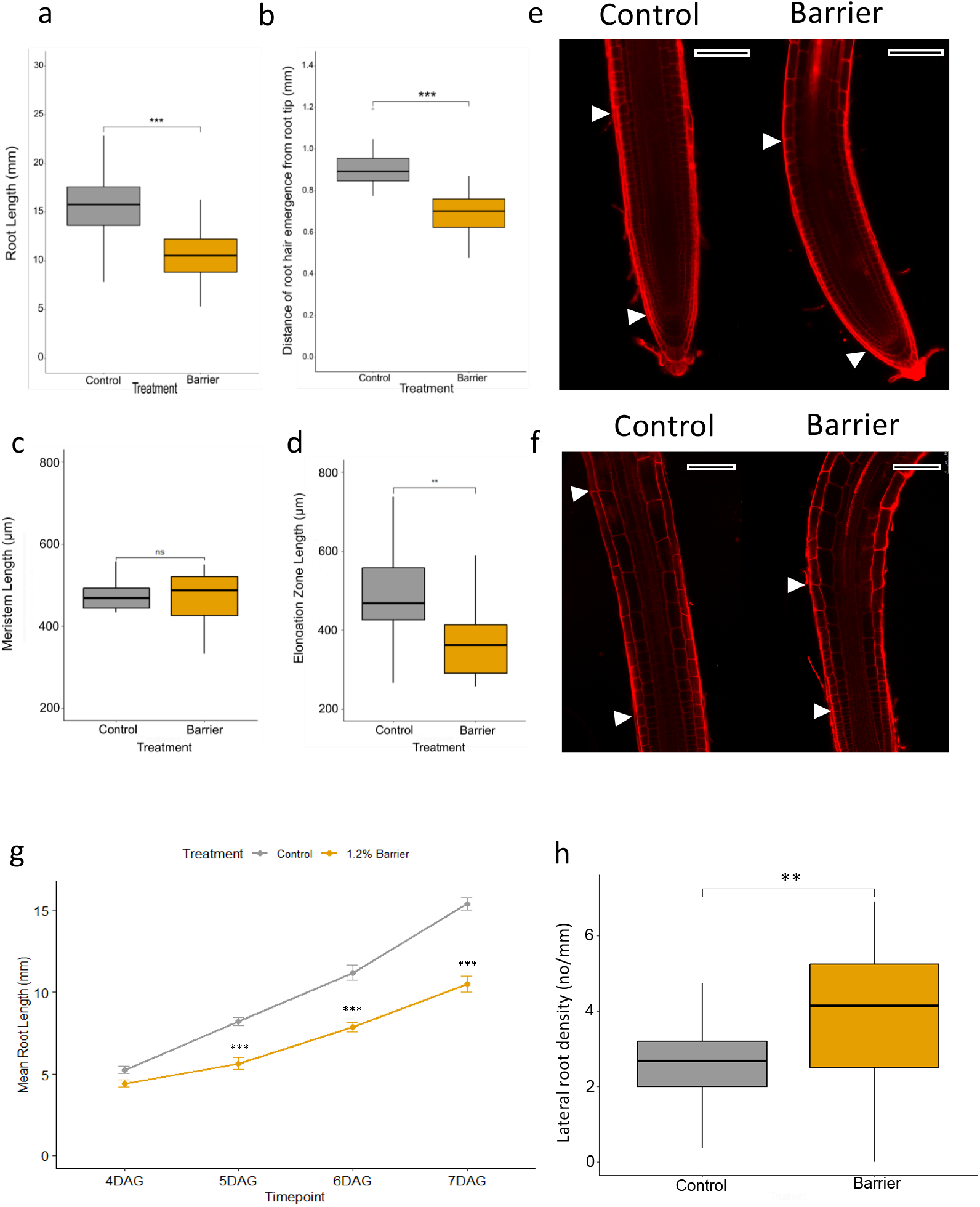
Barrier effects on primary root growth. Seeds were sown in a top layer of 0.3% Phytagel and roots grew for 5mm before encountering the lower media layer. a) Root length of 7 DAG seedlings (Student’s *t*-test, P < 0.001). b) Distance of root hair emergence from the root tip of 7 DAG seedlings (Student’s *t*-test, P < 0.001). c) Meristem length at 7 DAG (Student’s *t*-test, p = 0.79). d) Elongation zone length at 7 DAG (Student’s *t*-test, P = 0.003). e) Primary root tip stained with propidium iodide at 7 DAG. Arrowheads indicate quiescent centre and end of the meristematic zone. f) Elongation zone of primary roots stained with propidium iodide. Arrowheads indicate start and end of the elongation zone. Scale bar = 50 μm. g) Mean primary root length between 4-7 DAG when encountering a barrier. For boxplots, upper and lower boundaries of the box indicate the interquartile range (IQR), a black line within the box marks the median, and whiskers represent the min and max excluding outliers. Open circles represent outliers. ns = not significant, * < 0.05, ** < 0.01, *** < 0.001. Error bars indicate mean ± SE.

Analysis of CYCB1;2:GUS reporter expression, which marks the G2/M cell cycle transition (Colon-Carmona *et al*., 1999; Schnittger *et al*., 2002), showed the number of dividing cells remained unchanged in response to impedance (Fig. S2 a,b). The ratio of dividing cells on the ‘left’ and ‘right’ sides of the axial root plane was calculated using the formula exp(|log(left/right)|), which obviates the need to assign left or right sides. This ratio does not change in impeded roots (Fig. S2c), indicating cell division rate remains constant radially across the meristem. While primary root length was significantly reduced in impeded roots, lateral root density (i.e. number of laterals per length of primary root) increased (Fig. 1h), suggesting a redistribution of resource to lateral roots in response to the barrier.

To examine the short-term (0-24 h) impedance response, seedlings were grown on vertical plates on the gel surface and at 6 DAG plastic barriers were placed horizontally directly in front of the growing roots. Consistent with previous studies (Massa & Gilroy, 2003; Lee *et al*., 2020), roots were found to rapidly bend away from obstacles, forming a ‘step-like’ growth pattern where the root grows parallel to the obstacle while the root tip maintains contact with the barrier. It is hypothesized that the first bend represents a touch response to the barrier, leading to ‘exploration’ of the substrate, while the second bend downwards represents a gravitropic response. Root tip angle (RTA) was measured between the leading edge of the root tip and the surface of the barrier, i.e. at the initiation of the second bend (Fig. S3a; Fig. S1d); this provides a measurable ‘readout’ of the response to mechanical impedance. RTA changed from an average of 93.8° ± 2.20° (0 h) to 120° ±1.25° (180-480 mins; Fig. S3b), detectable within 60 min of the root encountering an obstacle, with a second bend forming between 3 and 4 hours. The rate of bending (ΔRTA/min) was greatest between 15-90 min (Fig. S3c); root growth rate was fairly constant along the barrier (average 0.96 ± 0.14 mm over 480 min; Fig. S3d).

After 6 h, meristem size was unaffected in roots responding to the barrier (Fig. S3e,f) consistent with changes in cell division unlikely to be involved in the early response to a barrier (Fig. S2; Okamato *et al*. 2008). However, the ratio of the length of the elongation zone between the two sides of the root significantly increased in response to a barrier, from a median ratio of 1.01 (IQR = 0.017) to 1.04 (IQR = 0.027), demonstrating asymmetry in cell elongation (Fig. S3g). The average ratio of cell length for the first 8-11 cells of the elongation zone significantly increased in response to a barrier, confirming the asymmetry (Fig. S3h).

### Transcriptional analysis reveals specific signalling changes in the impeded root

The results described show mechanical impedance induces reduced root growth and step-like bending due to differential cell elongation, rather than altered meristem activity, enhanced root hair growth and increased lateral root density. To understand better the molecular events in roots under mechanical impedance stress, time-course RNA-sequencing (RNA-seq) was used to identify transcriptional changes for 3 independent biological replicates for control and barrier treatments of 6 DAG roots at 6 h and 30 h after impedance, representing relatively early and late stages of ‘two-step’ growth responses.

Principal component analysis (Fig. S4a) and a heatmap of the 50 most highly expressed genes with clustering across samples (Fig. S4a) show that variation between sample groups is greater than variation within groups. A P-value of <0.05 and Log2 fold change (log^2^fc) of >0.5 or <-0.5 were selected to identify differentially expressed genes (DEGs) between barrier and control conditions at each time-point (Fig. S5a). 1941 genes were upregulated and 406 downregulated at 6 h after barrier placement. Fewer genes were differentially expressed at 30 h, with 852 upregulated and 607 downregulated genes (Fig. S5b). 372 genes were differentially expressed at both 6 h and 30 h (Fig. S5c).

In an hypothesis-building process, we carried out an unbiased gene ontology (GO) enrichment analysis to interpret the biological functions of genes upregulated and downregulated at each timepoint, and treemaps were generated using data from REVIGO (Supek *et al*., 2011). Results reveal little overlap between GO terms in the response at 6 h versus 30 h after impedance. Of particular interest, there is seen a strong upregulation in genes involved in ‘response to stress’, the largest supercluster of GO terms, at 6 h (Fig. S6). GO terms that relate to reactive oxygen species (ROS) are enriched, including ‘hydrogen peroxide catabolism’ and ‘reactive oxygen species metabolism’. Also identified was an enrichment in genes involved in ‘signalling’, ‘protein phosphorylation’, ‘cell communication, ‘secondary metabolism’ and ‘localization’. At 30 h, there is an upregulation of genes involved in mRNA splicing, cellular respiration and ribonucleoprotein complex biogenesis (Fig. S7). In contrast to the response at 6 h, genes involved in ‘cell communication’, ‘signalling’ and ‘response to stimulus’ are downregulated at 30 h (Fig. S7), indicating a switch from the response at 6 h.

At 6 h we found that GO terms relating to hormone responses were enriched in upregulated genes (Fig. S8). In contrast, the majority of genes relating to hormone signalling at 30 h are downregulated, consistent with the observation that many stress-related genes are also downregulated after 30 h (Fig. S8). This indicates that hormone-related transcriptional changes principally occur relatively early in the response to mechanical impedance. Although transcriptional upregulation of genes associated with GA and ABA signalling were identified, analysis of RGA:GFP expression (GA signalling) and treatments with fluoridon (for ABA signalling; Rowe *et al*., 2016) provided no further evidence for an essential role for these pathways in the early impedance response (data not shown); further analysis therefore focused on ROS, ethylene and auxin requirements.

### A role for ROS in the impedance response

ROS have been proposed to act as a rapid wave-like signal during stress responses, mediated by RESPIRATORY BURST OXIDASE HOMOLOG D (RBOHD) activation (Miller *et al*., 2009; Gilroy *et al*., 2014, 2016). GO analysis showed enrichment in terms relating to ROS metabolism. At 6 h there was an upregulation in genes relating to ‘reactive oxygen species metabolism’ and ‘hydrogen peroxide catabolism’. NADPH oxidases (respiratory burst oxidase homologues) are key ROS-producing enzymes and act as molecular ‘hubs’ during ROS-mediated signalling (Hu *et al*., 2020). Six NADPH oxidase genes are upregulated at 6 h (Table S1), including *RBHOD* and *RBOHF*, which have been shown to be involved in a number of abiotic and biotic stress responses (Xie *et al*., 2011; Liu *et al*., 2015; Morales *et al*., 2016; Wang *et al*., 2019). *ROOT HAIR DEFECTIVE 2* (*RHD2*) is upregulated at 6 h, and has been linked to root hair growth (Foreman *et al*., 2003) and root touch responses (Monshausen *et al*., 2009). In addition, 20 known ROS scavenging genes were upregulated at 6 h (Table S2).

Transcriptomic changes in response to ROS have previously been documented and a transcriptomic footprint created (Gadjev *et al*., 2006). Comparison of this footprint with the data obtained for response to a barrier revealed that, at 6 h, 32 upregulated genes overlap with the ROS footprint (Fig. 2a). The greatest number of DEGs at 6 h appear to be upregulated in response to singlet oxygen (Fig. 2b), strongly suggesting an elevation of ROS transcriptional and signalling responses when a root encounters a barrier.

**Figure 2.**
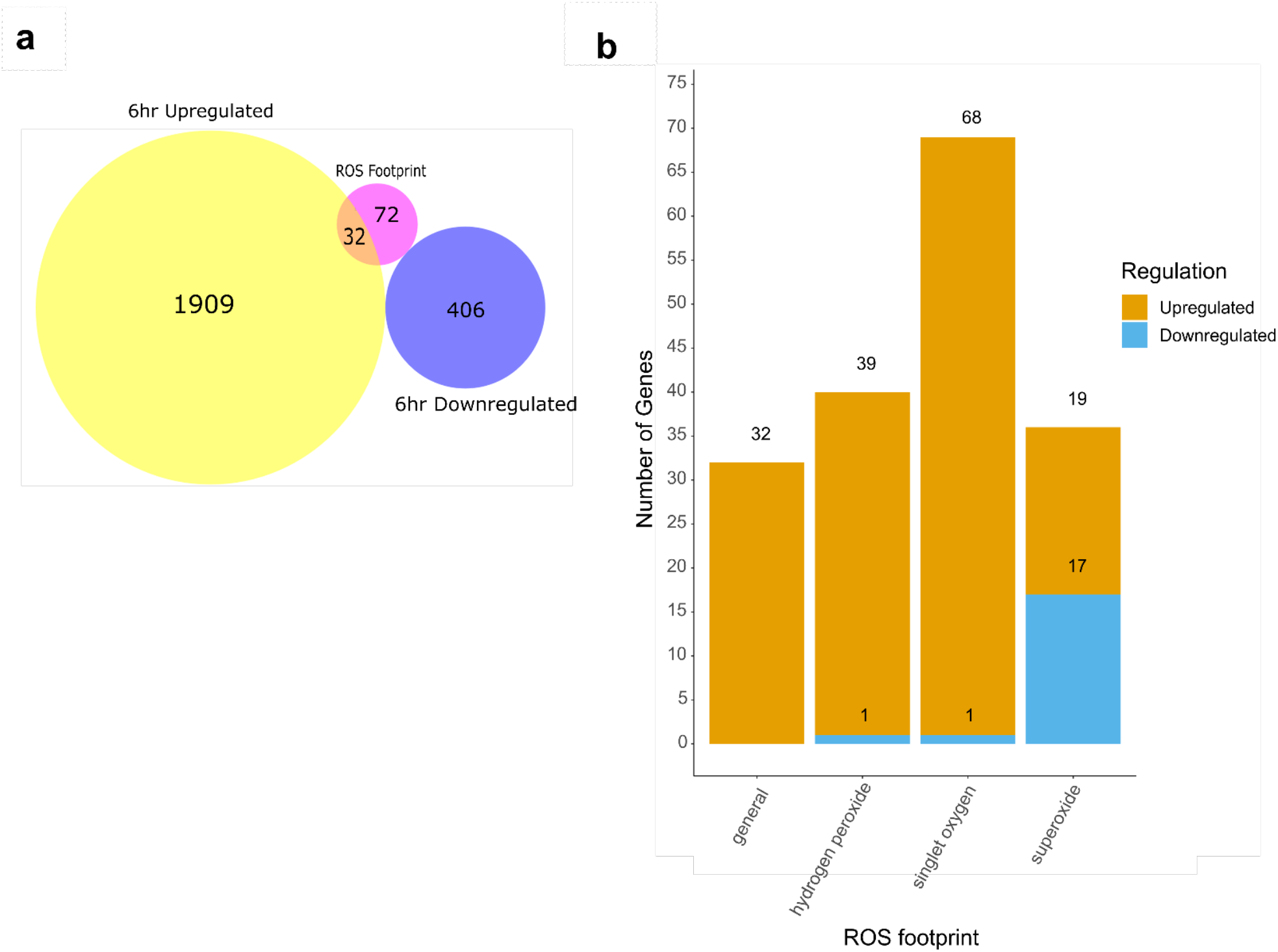
ROS-related gene expression at 6 h in response to a barrier. a) Venn diagram showing common genes between genes differentially expressed at 6 h in response to a barrier and genes identified as a general ROS transcriptomic footprint (Gadjev *et al*., 2006). b) Number of genes identified from the RNA-Seq data set at 6 h identified by Gadjev *et al*. (2006) as relating to specific reactive oxygen species responses.

Since *RBHOD* and *RBOHF* genes were upregulated within 6 h, time-lapse imaging was used to examine the early root response of the *atrbohD/F* double mutant after encountering a barrier (Fig. 3a). The *atrbohD/F* root tip angle differed significantly from Col-0 at 360 min, with mean RTA of *atrbohD/F* of 148.8° (± 5.7) compared with 118.9° (± 8.2) in the control (Fig. 3b; ANCOVA, p < 0.001); differences were detected from ca. 75 min. At 75 min the RTA of *atrbohD/F* is greater at 124.0° (± 6.7) compared with 108.9° (±4.4) for Col-0 (Fig. 3b), demonstrating a requirement for these ROS signalling pathway genes in the early impedance response.

**Figure 3.**
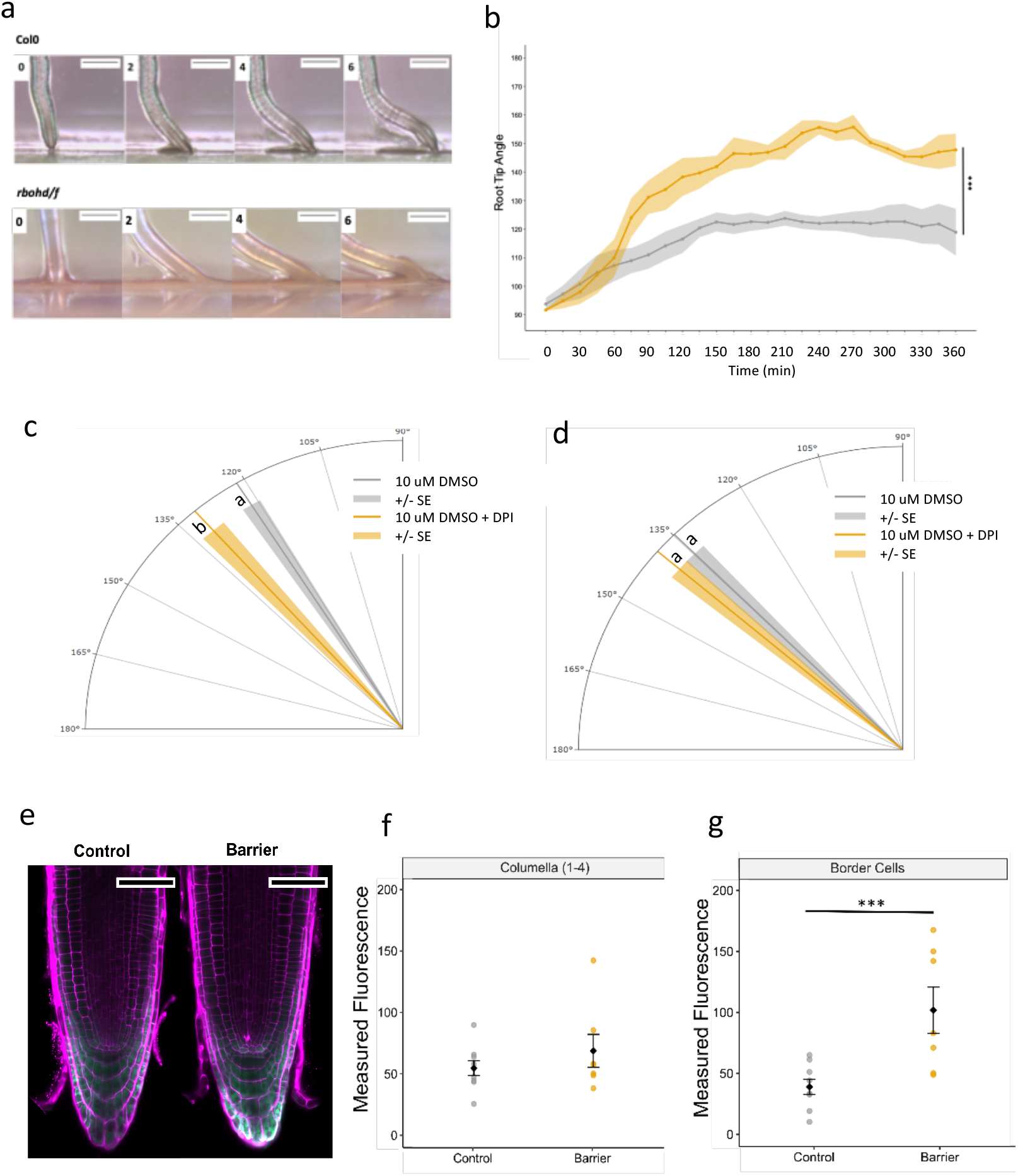
ROS is required for the barrier response. a) Timelapse imaging of *atrbohD/F* double mutant in response to a barrier. Plastic barriers were placed in front of 6 DAG vertically growing roots and root tips were imaged every 15 min. b) RTA from 0-360 min] after contact with a barrier. Root tip angle was measured every 15 min. c,d) The effect of diphenylene iodonium (DPI) on root tip angle during the barrier response 6 h after barrier placement. Lines indicate mean and surrounding shaded area indicates ± SE. e-g) HyPer fluorescence in unfixed roots after responding to a barrier. Barriers were placed in front of seedlings 6 DAG and roots were imaged between 3-5 h after barrier placement. Roots were stained with propidium iodide prior to imaging. e) Typical fluorescence of Hyper probe YFP excited at 488 nm (Green) in roots stained with propidium iodide (magenta). f) Measured fluorescence of Hyper YFP in the columella. g) Measured fluorescence of Hyper YFP in root cap border cells.

Diphenylene iodonium (DPI), a chemical inhibitor of ROS production, was used to investigate further the role of ROS in the barrier response. Seedlings were grown on 10 μM DPI or 10 μM DMSO as a control, and barrier response determined. At 6 h, seedlings grown on 10 μM of DPI encountering a barrier exhibited a higher RTA compared to the control (10 μm DMSO; Fig. 3c), confirming the mutant study. At 24 h, there was no significant difference in RTA between seedlings grown in the presence of 10 μM DPI on 10 μM DMSO when encountering a barrier (Fig. 3d).

Confocal imaging of the H_2_O_2_ reporter HyPer (Belousov *et al*., 2006) was used to investigate possible changes in H_2_O_2_ levels at the root tip in response to impedance within 6 h. YFP fluorescence increased significantly in the root cap border cells, indicating an increase in H_2_O_2_ (Fig. 3e,f). There was no significant change detected in other cells of the root cap (Fig. 3g,h).

CellROX Deep Red is a cell-permeant dye that is weakly fluorescent in the reduced state and exhibits photostable fluorescence upon oxidation by ROS. Three hours after barrier placement, fluorescence was observed in the outermost columella, border cells and in the lateral root cap (Fig. 4a) but was not significantly different in either the columella cells or lateral root cap in the two treatments (Fig. 4b) Fluorescence was also observed in both the meristematic and elongation zones (Fig. 4c), and while there was no difference in meristem cells, there was found a significant decrease in CellROX fluorescence in the elongation zone in roots encountering a barrier (Fig. 4d, Student’s *t*-test, p = 0.05), indicating a decrease in ROS.

**Figure 4.**
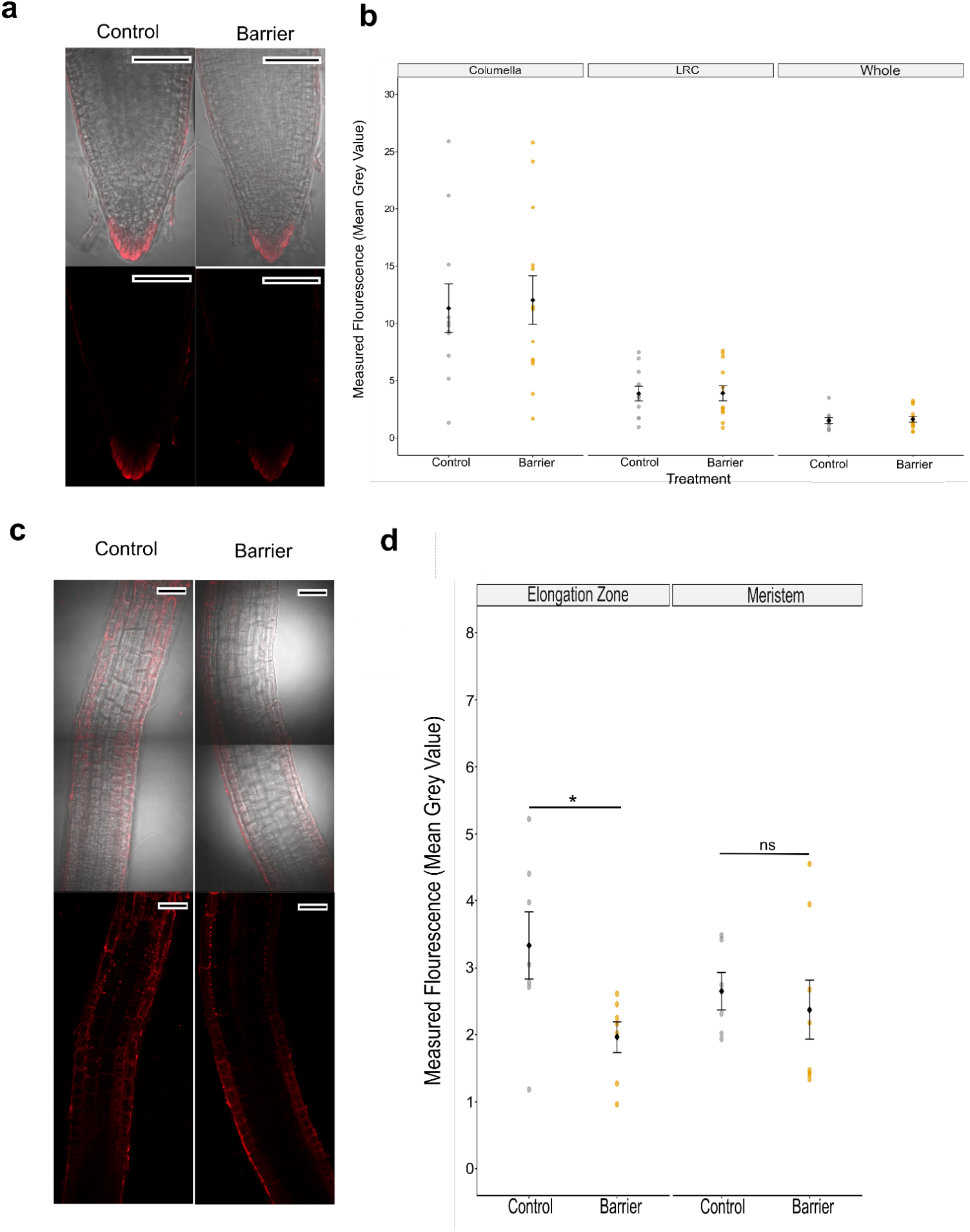
CellROX staining to reveal ROS levels. Barriers were placed in front of seedlings 6 DAG and seedlings were removed 3 h after barrier placement for staining with CellROX Deep Red. a) Typical staining pattern of CellROX Deep Red (Red) at the root tip. With and without bright field image of the root tip for reference. b) Measured fluorescence of CellROX Deep Red stain in the root tip. c) Typical staining pattern of CellROX Deep Red (Red) in the meristem and elongation zone, with (upper panels) and without (lower panels) bright field imaging. d) Measured fluorescence of CellROX Deep Red stain in the meristem and elongation zone. e) Ratio of CellROX Deep Red fluorescence across the left and right side of the meristem and elongation zone. Ratio was calculated using the formula exp(|(log(left/right))|) to account for any bias in assigning left/right side of the root. Scale bar = 50 μm. Black circles and error bars represent mean ± SE. Coloured circles represent distribution of individual data points. Lines and asterisks show significance (ns = not significant, * < 0.05, ** < 0.01, *** < 0.001).

### A role for ethylene in the impedance response

KEGG pathway mapping identified upregulation of ethylene-related transcription at 6 h, including key ethylene response genes such as *ETHYLENE INSENSTIVE LIKE 2 (EIL2;* Fig. S9a) and two *ETHYLENE RESPONSE FACTOR* (*ERF*) genes *ERF1A* and *ERF4*, and genes involved in ethylene biosynthesis - *ACS5* is upregulated by log^2^FC of 2.52, and the ACC oxidase gene *ETHYLENE FORMING ENZYME* (*EFE*) is also upregulated at 6 h, suggesting increased ethylene production in response to a barrier (Fig. S9b). At 30 h, *ACS5* and *ELO1* are both downregulated, indicating a possible decrease in ethylene biosynthesis. These data suggest that upregulation in ethylene biosynthesis may be early and transient during the barrier response.

Previous studies describe controversy around a role for ethylene synthesis, and suggested primarily a role for ethylene signalling, in the response to continuous impedance (Okamoto *et al*., 2003). To investigate a possible requirement for ethylene signalling for the root response, time lapse imaging was used to investigate the bending of the ethylene resistant *etr1-1* mutant in response to a barrier (Fig. 5a). RTA changed more rapidly in *etr1*compared with Col-0 in response to a barrier, detectable from ca. 45-60 min, indicating a relatively early role of ethylene signalling in the impedance response. At 60 min, mean RTA of *etr1* was 122.9° ± 3.40, 15.6 degrees higher than the root tip angle of Col-0 (107.3° ± 5.53). The RTA of *etr1* also reduced again toward the vertical between 195 and 270 min, reaching 117.6° ±9.9 at 270 min before increasing again to 150° ±12.9 at 360 min (Fig. 5b,c), associated with the second bend. *ein2* also showed a slightly more vertical root than wildtype (though not significant at the P<0.05 level; Fig. S10), indicating a role for ethylene signalling in reducing growth rate and bending on impedance. *etr1* seedlings grew longer than wildtype in a gel barrier (Fig. S11) and roots of a range of ethylene-insensitive mutants (*ein2, etr1, aux1, eir1*) were not significantly inhibited by barrier impedance (ANOVA, p<0.0001; Fig. S12).

**Figure 5.**
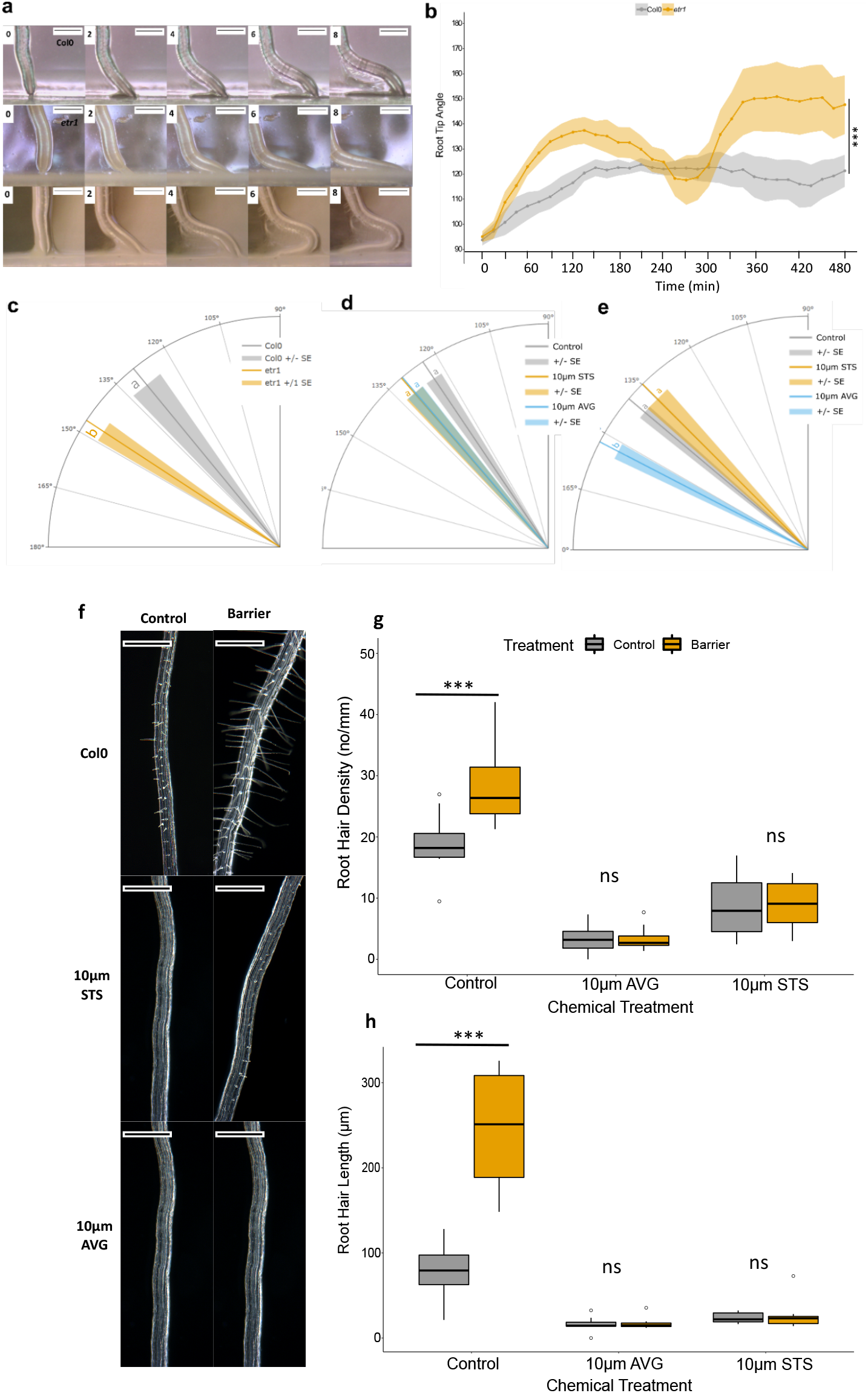
Ethylene signalling is required for the barrier response. a) Timelapse imaging of wildtype (Col-0, upper panels) and *etr1* (lower panels) between 0 and 8 h after barrier placement. Plastic barriers were placed in front of 6 DAG vertically growing roots and root tips were imaged every 15 minutes. In some cases, *etr1* shows a reversal of growth direction (lower panels at 6 and 8 h). b) RTA from 0-480 minutes after contact with a barrier. Root tip angle was measured every 15 min. The central ‘dip’ in *etr1* RTA reflects the directional change. Lines and dots indicate mean with shaded area indicating ± SE. Scale bar = 0.5 mm. c) Angle of primary root tips to the horizontal barrier 24 h after barrier placement for wildtype (Col-o) and *etr1*. d,e) Effect of AVG and STS on RTA 6 h (d) and 30 h (e) after barrier placement. Lines indicate mean and surrounding shaded area indicates ± SE. f) Typical root hair growth after 24 hours in the presence or absence of a barrier: untreated (upper panel, in presence of STS (middle panel), in presence of AVG (lower panel). Scale bar represents 500 μm g) Root hair density (number of root hairs/mm root). Root hairs were counted in an approximate 2 mm section of mature root and exact distance measured to calculate root density. h) Root hair length. Root hair length was measured in 20 root hairs within a 2 mm region of root and the average taken to determine root hair length for the sample. Brackets and asterisks indicate significance with a Tukey Pairwise comparison (ns = not significant, * < 0.05, ** < 0.01, *** < 0.001).

To further examine the role of ethylene, seedlings were grown in the presence of chemical inhibitors of ethylene biosynthesis (10 μM AVG) and signalling (10 μM silver thiosulphate, STS). At 6 h after barrier placement, seedlings grown in the presence of either inhibitor showed a slightly higher RTA than the control, however this was non-significant (Fig. 5d, ANOVA, p = 0.08). After 24 h of growth, seedlings grown on 10 μM AVG had a significantly higher mean RTA than the control (Fig. 5e; ANOVA, p < 0.001; TUKEY p = 0.003), indicating a role for ethylene synthesis at the later stages of impedance response (likely associated with gravitropism). However, seedlings grown in the presence of STS did not exhibit any significant change in RTA compared with the control (Fig. 5e, TUKEY, p > 0.05).

Root hair density and length were measured in roots responding to a barrier in the presence of 10 μM AVG or STS. At 24 h of barrier treatment, control seedlings showed an increase in root hair length in response to a barrier as described above, which together with root hair density was significantly reduced by treatment with 10 μM AVG or 10 μM STS in either the presence or absence of barrier (Fig. 5f-h; ANOVA, P <0.001). This shows that normal ethylene biosynthesis and signalling are each essential for root hair elongation and formation, but only root hair length increases in response to barrier contact.

### A role for auxin in the impedance response

Further KEGG pathway mapping revealed DEGs associated with auxin signalling. At 6 h *PHYTOCHROME-ASSOCIATED PROTEIN 2* (*PAP2*), a member of the *AUXIN/INDOLE-3-ACETIC ACID* (*Aux/IAA*) family, and *SAUR36* are downregulated, while *SAUR55* is upregulated; *SAUR36* is upregulated at 30 h. At 30 h, 3 *AUX/IAA* genes are differentially expressed: *IAA30* and *PAP2* are upregulated while *IAA14* is downregulated (Fig. S13). Three genes involved in auxin conjugation, *GH3.17, BRU6* and *DFL1* (Staswick *et al*., 2005) are also upregulated at 6 h and two downregulated at 30 h (*GH3.17* and *DFL2*, Fig. S13), suggesting a role for auxin conjugation in controlling free auxin levels during the barrier response. Three ATP-Binding Cassette (ABC) family genes, *ABCB1, ABCB4* and *ABCG37*, associated with auxin transport and gravitropic response (Geisler *et al*., 2005; Wu *et al*., 2007; Lewis *et al*., 2007; Růžička *et al*., 2010) are upregulated at 6 h.

The transcriptomic data presented above, and recent work by Lee *et al*. (2020), support a role for auxin and its transport in obstacle avoidance. Using time lapse imaging, we found that RTA for the auxin transport (and ethylene-insensitive) mutants *eir1-4* and *aux1-7* was significantly closer to the horizontal than Col-0 (Fig. 6a,b), apparent from ca. 210 min (average RTA for *eir1-4* is 133 ± 5.5° and 138.4 ± 3.4° for *aux1-7*, compared with 123.8 ± 2.6° for Col-0). Initial bending between 0 and 210 minutes appears the same between Col-0 and the mutants (Fig. 6b). After 24 h the RTA was still significantly different between Col-0 and the mutants (ANOVA, P <0.001; Fig. 6c). Wildtype seedlings treated with the auxin transport inhibitor N-1-naphthylphthalamic acid (10 μM NPA) at 6 h after barrier interaction had a significantly higher RTA than untreated controls (161 ± 4.4°) (Fig. 6d) and at 24 h mean RTA at both 2.5 and 10 μM NPA was greater than the untreated controls (Fig. 6e).

**Figure 6.**
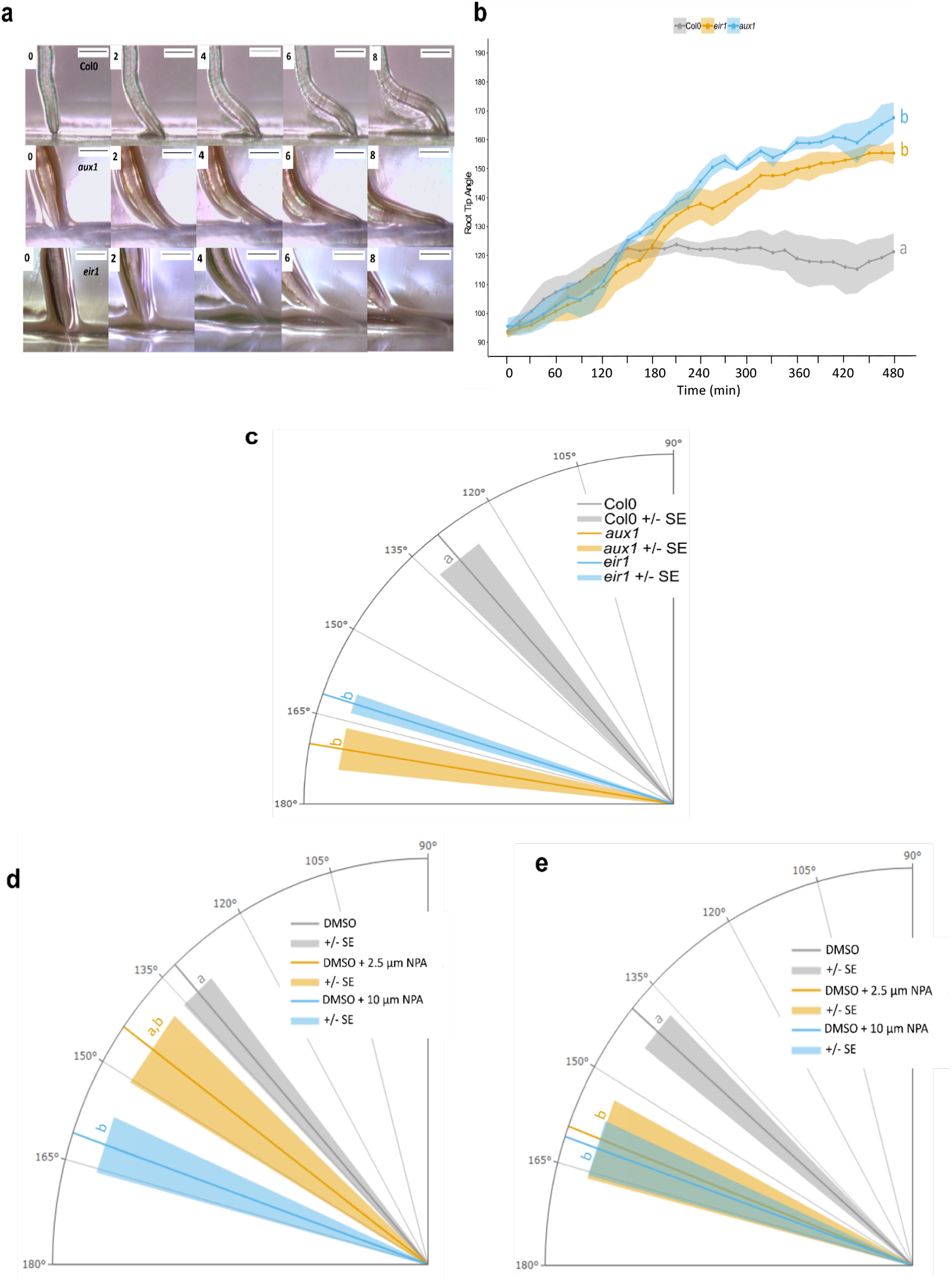
Ethylene signalling is required for the barrier response. a) Timelapse imaging of wildtype. (Col-0), *aux1* and *eir1* between 0 and 8 h after barrier placement. Plastic barriers were placed in front of 6 DAG growing roots and root tips were imaged every 15 min. b) RTA over time from 0-480 minutes after contact with a barrier. Root tip angle was measured every 15 minutes. Lines and dots indicate mean with shaded area indicating ± SE c) Angle of primary root tips to the horizontal barrier 24 h after barrier placement. d,e) Effect of NPA on RTA during the barrier response 6 h (d) or 24 h (e) after barrier placement. Lines indicate mean and surrounding shaded area indicates ± SE Scale bar = 0.5 mm. Letters indicate significance with a Tukey Pairwise comparison, P < 0.05.

Mechanical impedance caused a significant increase in fluorescence of the reporter line DR5rev::3xVENUS-N7 tip 6 h after encountering a barrier in the stele and lateral root cap (Fig. 7a,b). Similarly, the ratiometric reporter R2D2 showed Tomato/Venus fluorescence increased significantly at 6 h (Fig. 7c-e), indicating an increase in auxin levels. Asymmetric distribution of auxin was also detected, with ‘left to right’ auxin level increasing significantly in response to impedance (Fig. 7d). There was also a significant increase in Tomato/Venus fluorescence by 4 h, i.e. the time when auxin transport mutants showed altered RTA responses (ANOVA p = 0.001, Fig. S14), indicating a significant increase in auxin at the root tip in that early response period. Cell type analysis also revealed a significant change in left/right auxin level in the LRC/Epidermis between 2 h and 4 h (ANOVA, p = 0.03; Fig. S14). These data show auxin distribution dynamics across the root occurring at early time points from the first bending impedance response.

**Figure 7.**
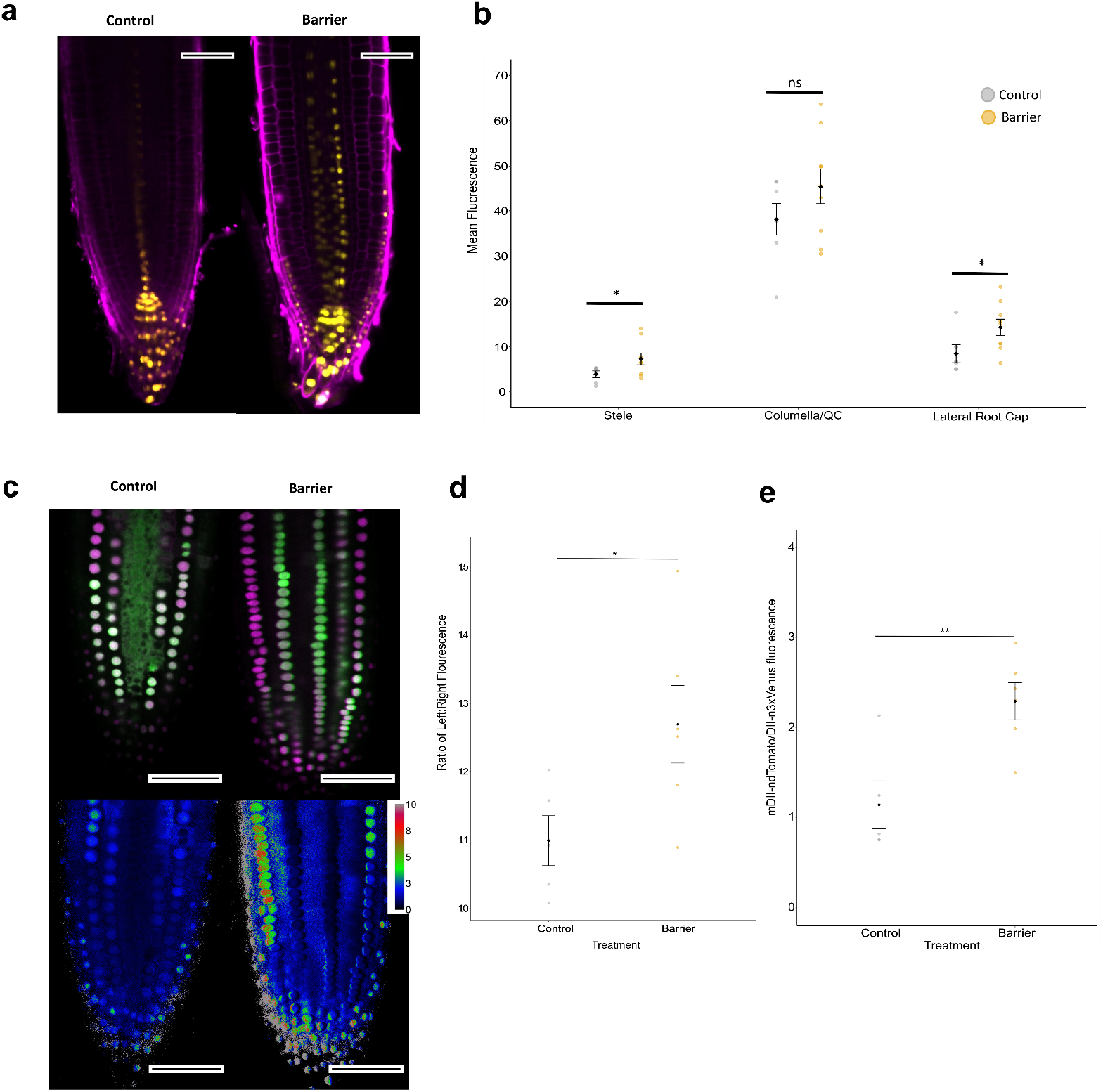
Imaging of reporters reveals auxin distribution changes. Barriers were placed in front of vertically growing roots 6 days after germination (DAG). a) Confocal imaging of the DR5rev::3xVENUS-N7 auxin reporter 6 h after placement of a barrier and stained with propidium iodide before imaging, compared with control. Yellow = Venus, Magenta = propidium iodide. b) Relative mean fluorescence measured using imageJ in the Stele, Columella/Quiescent Centre (QC) and Lateral Root Cap (LRC). c) Confocal imaging of the R2D2 auxin reporter in roots responding to a barrier at 6 h. Green = DII-m3xVenus, Magenta = mDII-ndTomato. Ratiometric image of mDII-ndTomato/ DII-m3xVenus fluorescence produced in imageJ using image calculator. d) Ratio of mDII-ndTomato/ DII-m3xVenus fluorescence. e) Ratio of auxin level across the left and right sides of the root tip. Ratio was calculated using the formula exp(|(log(left/right))|) to account for any bias in assigning left/right side of the root. Scale bar = 50 μm. Black circles and error bars represent mean ± SE. Coloured circles represent distribution of individual data points. Lines and asterisks show significance (ns = not significant, * < 0.05, ** < 0.01, *** < 0.001).

## Discussion

Previous studies have investigated the response of roots to mechanical stimulation through the use of artificial obstacles to growth or changes in orientation of growing roots. Alterations in root system morphology are observed when a plant is reoriented with respect to the direction of the gravitational field. For example, *Arabidopsis* roots produce a characteristic wavy growth pattern when grown on tilted agar plates with impenetrable surfaces (Okada & Shimura, 1990; Rutherford & Masson, 1996). These patterns of growth are the result of a combination of gravity sensing and touch stimulation from the agar surface (Oliva & Dunand, 2007).

Placing a horizontal obstacle in the way of a vertically growing root reveals another pattern of growth likely to be induced by a combination of touch sensing and gravity, the barrier response. When encountering a barrier, *Arabidopsis* roots show a ‘step-like’ growth, with only the root tip remaining in contact with the barrier surface (Massa & Gilroy, 2003; Lee et al., 2020). Root bending has also been observed in roots grown in a medium consisting of a soft upper layer and a hard lower layer. The root exhibits a bending response at the lower, harder layer (Yamamoto *et al*., 2008; Yan *et al*., 2017, 2018). It has been suggested that a zone of ‘mechanical weakness’ is required for the bending process and that this is localised between the growing and mature zones of the root (Bizet *et al*., 2016).

While the tropic responses of roots have been characterized in different substrates with different mechanical barrier properties, there is limited information on the molecular responses and controlling mechanisms. The first bend, on touching the barrier, is likely a touch response mediated, it has been suggested, by cells of the root cap (Massa & Gilroy, 2003) followed by a gravitropic response to keep the root tip pointing downwards while the elongation of the root causes lateral movement; this response appears to require polar auxin transport via PIN2 (Lee *et al*., 2020).

In our system, we similarly observed the two-step growth pattern, and the RNA-seq analysis revealed significant changes in three, likely inter-related, signalling pathways, namely ROS, ethylene and auxin. From an analysis of the RTA of ROS, ethylene signalling and auxin transport mutants and reporter imaging studies, it would appear that ROS and ethylene responses occur relatively early, during the period of the touch and first bend of the root, followed by auxin responses. The root behaviour of the *atrbohD/F* double mutant, defective in ROS generation, is first detectable within ca. 75 min of barrier touching, followed by Hyper and CellROX ROS reporter accumulation in root cap border cells and lateral root cap and root elongation zone within 3 h (Figs. 3 & 4). No significant barrier effect on ROS reporter activity was seen in the root meristem. Early ethylene requirements were also seen as altered root behaviour of the ethylene-insensitive *etr1* mutant within 45-60 min of barrier contact. Over a longer period (6-24 h), a likely role for ethylene biosynthesis was identified through root responses of seedlings (RTA, root hair density and length) treated with the ethylene synthesis inhibitor AVG (Fig. 5). However, our RNA-seq data suggest an upregulation in ethylene signalling and biosynthesis at the transcriptional level is transient. At 30 h, *ACS5* is downregulated and there is no upregulation of any ethylene signalling genes. When ethylene signalling is perturbed, decreases in root growth in response to a barrier are not observed (Okamoto *et al*., 2008; Okamoto & Takahashi, 2019). The Arabidopsis ethylene-insensitive mutants either grow longer or are less inhibited than wildtype following barrier contact (Figs. S11, S12), suggesting that ethylene signalling may represent a useful target for breeders hoping to improve root growth in compact soils.

Consistent with recent previous work (Lee *et al*., 2020), auxin transport effects were observed through altered bending responses of *aux1* and *eir1* (*pin2*) mutants (which also exhibit ethylene sensitivity) or seedlings treated with the auxin transport inhibitor NPA, but not until after ca. 6-24 h after barrier contact; although auxin-related transcriptional changes were established by 6 h (Fig. 6).

Plant signalling systems act in an interacting network to elicit developmental change, such as in root growth and developmental responses to environmental stresses in the soil (Moore *et al*., 2015; Rowe *et al*., 2016). It is known for example that ethylene can lead to reduced primary root growth in Arabidopsis by activating auxin biosynthesis in the root tip and promoting its transport to the elongation zone, where it inhibits cell elongation (Ruzicka *et al*., 2007; Swarup *et al*., 2007). Given this evidence, and the observed timing of ethylene and auxin responses on contact with a mechanical barrier, it seems likely that ethylene signalling and/or synthesis changes lead to altered root growth and bending by effects on auxin distribution, as seen in Figs. 7 & 8. It is also likely that PIN2 is involved in the observed auxin redistribution (Lee *et al*. 2020), though other PIN proteins, AUX1, and possibly ABC transporters may also be involved. The activation of root hair development in response to barrier contact may represent a mechanism to anchor the root in the soil to facilitate the observed lateral growth around the barrier, as part of the soil exploration process.

**Figure 8.**
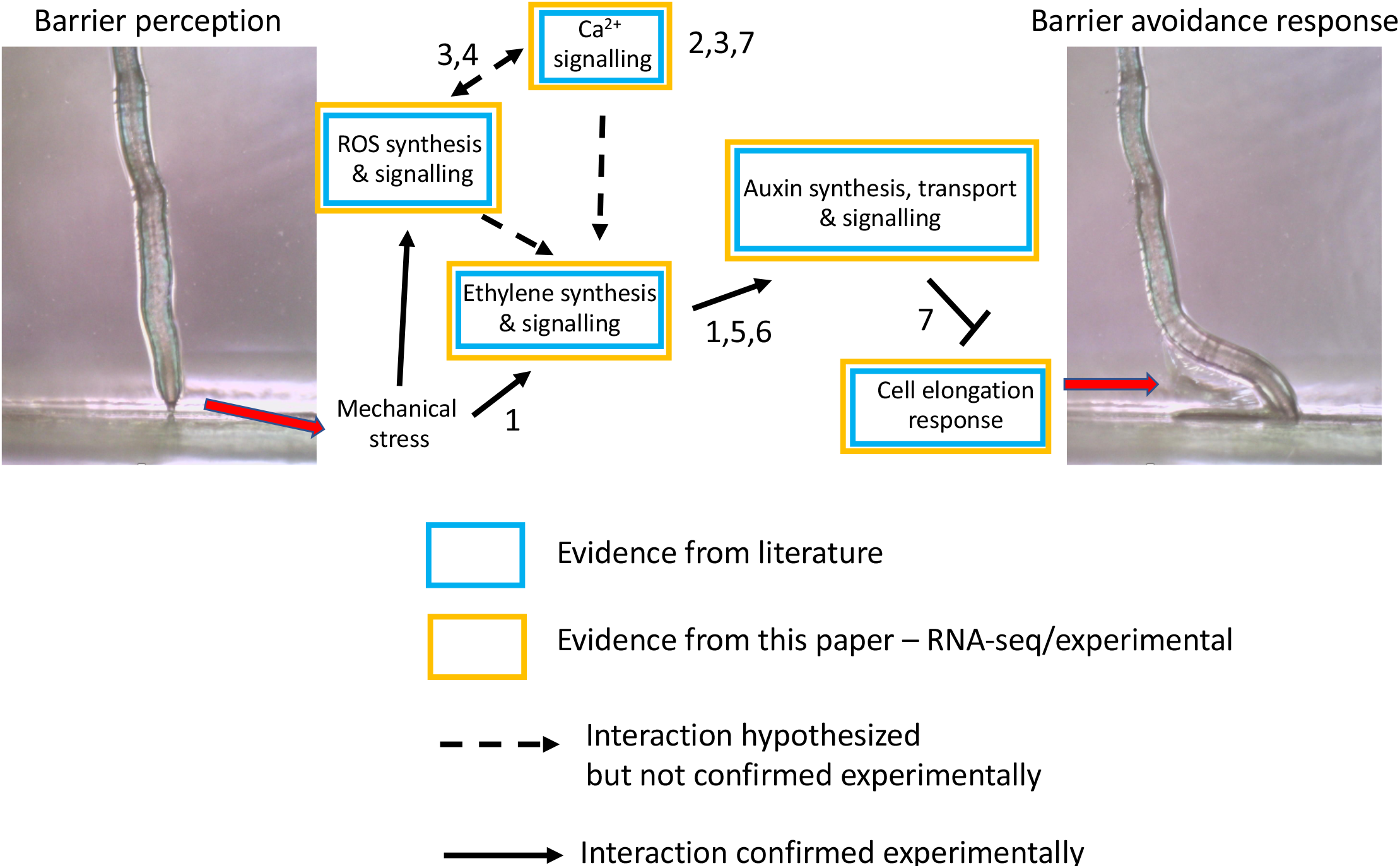
Pathways involved in the root barrier response. Hypotheses generated from results presented throughout the thesis and from previously published literature (numbers). Arrows show hypothesised positive interaction/relationship; t-bars inhibition or negative relationship. Boxed are colour coded according to the source of evidence. Numbers refer to published literature 1. Okamoto *et al*. (2008); Okamoto & Takahashi (2019) 2. Shih *et al*. (2014) 3. Monshausen *et al*. (2009) 4. Gilroy *et al*. (2014, 2016) 5. Růžička *et al*. (2007) 6. Strader *et al*. (2010) 7. Lee *et al*. (2020).

An interesting question is around the role of ROS. The evidence presented shows a relatively rapid upregulation of ROS-related gene transcription, including of genes encoding ROS-generating enzymes such as RHD2, required for root hair development and other stress responses (Foreman *et al*., 2003). What is unclear at present is the mechanistic relationship between ROS production and the activity of other signalling pathways, such as ethylene and auxin (Fig. 8). Inhibition of ROS signalling, such as in the *atrbohD/F* mutant or following treatment with the ROS inhibitor DPI, led to early changes in the root response to mechanical impedance, but how this might in turn lead to an activation of ethylene signalling (which in turn can induce auxin synthesis and transport) remains to be discovered. Monshausen *et al*. (2009) have demonstrated that ROS production in response to mechanical stimuli requires Ca^2+^ signalling. ROS and Ca^2+^ have been proposed to act together, with NADPH oxidase-produced ROS activating Ca^2+^ channels and the increase in Ca^2+^, in turn further activating NADPH oxidase activity (Gilroy *et al*., 2016). Recently, Wu *et al*. (2020) identified the first H_2_O_2_ receptor in plants, HPCA1. This membrane bound receptor kinase was also shown to mediate H_2_O_2_-induced activation of Ca^2+^ channels in guard cells, providing further evidence for how ROS mediates Ca^2+^ signalling. Data from our RNA-seq at 6 h after barrier contact did reveal an upregulation of this gene, three related genes (*HPCAL1, HPCAL2, HPCAL4*) and also genes involved in Ca^2+^ signalling, binding and transport, suggesting changes in levels of Ca^2+^ in the root. Future work should aim to explore further the interactions between ROS and Ca^2+^, and auxin and ethylene signalling during the root barrier response.

## Acknowledgements

KL acknowledges The Biotechnology and Biological Sciences Research Council (BBS/B/0773X) and Durham University for funding.

## Author contributions

KL and JFT devised the project; AGRJ carried out the experimental work; KL, JFT, and JX supervised the work; KL and AGRJ drafted the manuscript; all authors edited the manuscript.

## Data availability statement

All materials and data described in this paper are available to readers from the corresponding author, upon reasonable request.

## Supporting Information

**Figure S1. Barrier systems.** a) Magenta pot split layer assay. b) Plastic barrier assay. c) Dialysis membrane barrier assay. d) Root tip angle.

**Figure S2. proCYCB1;2::CYCB1:2:GUS expression reveals cell division in impeded roots.** a) Typical GUS staining pattern of CYCB1;2:GUS in roots grown in either a single layer of Phytagel (control) or a split layer system consisting of a lower, harder layer to impede growth. b) Number of dividing cells in presence of absence of barrier. c) Ratio of number of dividing cells between the left and right side of the meristem in presence of absence of barrier. Ratio was calculated as exp(|(log(left/right))|). For boxplots, upper and lower boundaries of the box indicate the interquartile range (IQR), a black line within the box marks the median, and whiskers represent the min and max excluding outliers. Open circles represent outliers. Asterisks and brackets indicate significance (ns = not significant, * < 0.05, ** < 0.01). Images and measurement representative of at least 15 samples. Scale bars = 50 μm.

**Figure S3. Short-term bending response of a root to a barrier.** Plastic barriers were placed in front of 6 DAG growing roots and root tips were imaged every 15 min. a) Time-lapse images of a Col-0 root tip encountering a horizontal barrier at 0-8 h after contact with the barrier. b) Root Tip Angle (RTA) over 0-480 min after contact with a barrier. RTA was measured every 15 minutes. c) Bending rate between 15-480 min. d) Root growth of the primary root tip between 0-480 min after contact with the barrier. Blue line represents Regression line. e) Typical elongation zone of primary root tips stained with Calcofluor White and grown in the presence or absence of a barrier for 6 h. Cells of the elongation zone used for measurements are outlined and highlighted in orange. f) Total cortical cell number in the meristem and elongation zone after 6 h. g) Ratio of meristem and elongation zone length between the left and right side of the root tip. h) Ratio of cell length between the left and right side of the root tip in the first 11 cells of the elongation zone. The root tip was divided through the middle into a left and right side and measurements taken separately for each side. Ratio was calculated using the formula exp(|log(left/right)|) to account for any bias in assigning left/right. Error bars represent mean ± SE. Scale bar = 50 μm.

**Figure S4. Analysis of variation in gene expression between and within sample groups.** a) Principal component analysis (PCA) plot visualising sample-sample distances. Polygons represent sample groups. PCA performed on regularized logarithm (rlog) transformed data using the R software package DEseq2. b) Clustered Heatmap of rlog transformed count data for the top 50 most highly expressed genes across all samples. Rows and column aggregated using kmeans clustering by the R package pheatmap.

**Figure S5. Genes differentially expressed in roots encountering a barrier compared with controls after 6 h and 30 h treatment identified through RNA-Seq.** a) Differentially Expressed Genes (DEGs) as estimated by the R software package DEseq2. Red dots represent significant DEGs, black dots non-significant. P-value < 0.05 and a log2 fold change (log2fc) > 0.5 or < - 0.5. b) Significant DEGs separated into up and downregulated genes at the two barrier response timepoints. c) Venn diagram representing overlapping DEGs between treatments.

**Figure S6.** Treemap output from REVIGO (Supek *et al*., 2011) of genes identified as significantly upregulated (a) and downregulated (b) 6 h after encountering a barrier. P-value < 0.05 and a log2 fold change (log2fc) > 0.5 or < −0.5. Each rectangle represents a gene ontology (GO) term cluster and each colour represents a supercluster of related clusters. Sizes of rectangles reflect the −log10 P-value of each cluster.

**Figure S7.** Treemap output from REVIGO (Supek *et al*., 2011) of genes identified as significantly upregulated **(a)** and downregulated **(b)** 30 hours after encountering a barrier. P-value < 0.05 and a log2 fold change (log2fc) > 0.5 or < −0.5. Each rectangle represents a gene ontology (GO) term cluster and each colour represents a supercluster of related clusters. Sizes of rectangles reflect the −log10 P-value of each cluster.

**Figure S8. Hormone signalling and metabolic/biosynthesis related GO terms identified by GO analysis of genes differentially expressed in response to a barrier**. Bar chart showing numbers of DEGs identified by each GO term. Bars are grouped by the hormone the GO term relates to.

**Figure S9. Ethylene-related gene expression analysis.** a) KEGG Pathway mapping of genes differentially expressed at 6 hours in response to a barrier and identified as being involved in the ethylene signalling pathway. Genes present within the data set are highlighted with colours corresponding to KEGG Orthology (molecular function) definition. b) Genes differentially expressed at 6 hours in response to a barrier and identified as being involved in the ethylene biosynthesis pathway. Components of the pathway with an identified DEG are highlighted red (ACC Synthase), orange (ACC Oxidase) and yellow (ETO1/EOL1).

**Figure S10. Response of *etr1* and *ein2* to a barrier.** a) Plastic barriers were placed in front of primary roots of seedlings 6 days after germination (DAG) and root tips were imaged at 24 hours after encountering a barrier. Scale bar indicates 0.5 mm. b) Angle of primary root tips to the horizontal barrier. Lines indicate mean and surrounding shaded area indicates ± SE. Letters indicate significance with a Tukey Pairwise comparison P < 0.05.

**Figure S11. Growth of *etr1* between 0 and 8 hours after barrier placement.** Root growth as measured by time-lapse imaging of roots encountering a barrier. a) Root growth of the primary root tip between 0-480 min after contact with the barrier. b) Root growth rate (μm/min) between 15-480 min after encountering a barrier. Plastic barriers were placed in front of 6 day old vertically growing roots and root tips were imaged every 15 min. Lines and dots indicate mean with shaded area indicating ± SE. Letters indicate significance after a *Student’s* t-test (*** < 0.001).

**Figure S12. Growth of wildtype and ethylene-sensitive mutant roots after barrier placement.** Primary root length was measured after growth of seedlings on dialysis membrane barriers for 7 days post germination. Error bars represent standard error of the mean of 15 seedlings per treatment. Root growth of the mutants was not significantly inhibited by barrier impedance (ANOVA and Tukey pairwise comparison, *** = P <0.0001, ** = P <0.001).

**Figure S13. KEGG Pathway mapping of genes differentially expressed at 6 and 30 hours in response to a barrier and identified as being involved in the auxin signalling pathway.** Genes present within the data set are highlighted with colours corresponding to KEGG Orthology (molecular function) definition.

**Figure S14. Confocal imaging of R2D2 in roots responding to a barrier between 0-4 h**. Barriers were placed in front of root tips 6 DAG. a) R2D2 fluorescence at the root tip at 0-4 h with accompanying ratio metric image of generated from mDII-ndTomato/ DII-m3xVenus fluorescence using ImageJ. For ratiometric images calibration bar indicates mean grey value. b) Ratio of mDII-ndTomato/ DII-m3xVenus fluorescence. c) Ratio of auxin level across the left and right sides of the root tip. Scale bar = 50 μm. Black circles and error bars represent mean ± SE. Coloured circles represent distribution of individual data points. Letters indicate significant difference after post-hoc TUKEY test, P < 0.05.

**Supplemental Table 1.** NADPH-oxidase genes identified through RNA-Seq that are upregulated during the barrier response at 6 hours.

**Supplemental Table 2.** List of genes that act as reactive oxygen species (ROS) scavengers identified in the RNA-Seq data of differentially expressed genes in response to a barrier log2fc identified with P-value <0.05.

