## Supplementary material for "Root growth responses to mechanical impedance are regulated by a network of ROS, ethylene and auxin signalling in Arabidopsis": Supplmental Figs and Tables

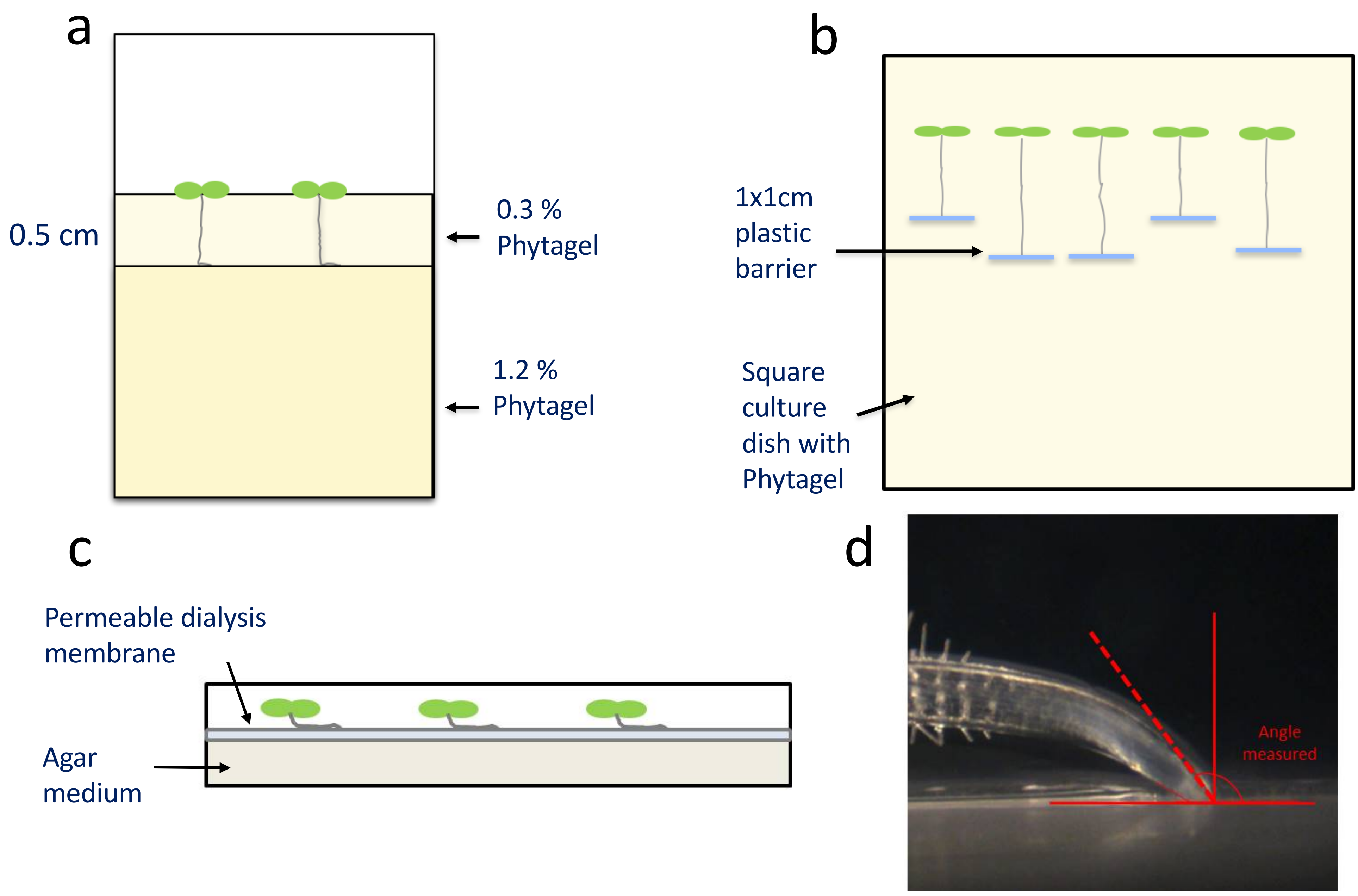


**Figure S1. Barrier systems.** a) Magenta pot split layer assay. b) Plastic barrier assay. c) Dialysis membrane barrier assay. d) Root tip angle.

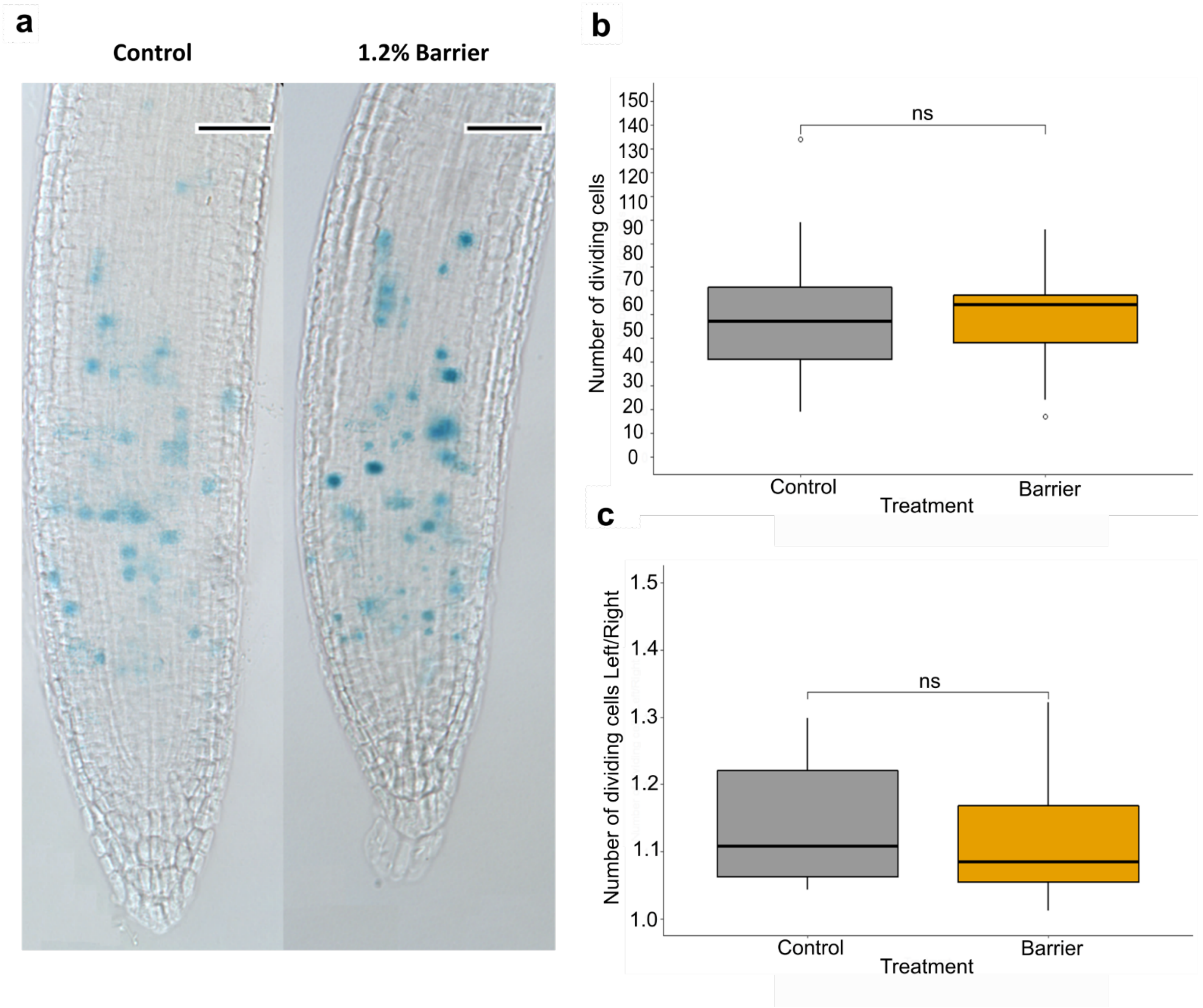


**Figure S2. proCYCB1;2::CYCB1:2:GUS expression reveals cell division in impeded roots.** a) Typical GUS staining pattern of CYCB1;2:GUS in roots grown in either a single layer of Phytagel (control) or a split layer system consisting of a lower, harder layer to impede growth. b) Number of dividing cells in presence of absence of barrier. c) Ratio of number of dividing cells between the left and right side of the meristem in presence of absence of barrier. Ratio was calculated as exp(|(log(left/right))|). For boxplots, upper and lower boundaries of the box indicate the interquartile range (IQR), a black line within the box marks the median, and whiskers represent the min and max excluding outliers. Open circles represent outliers. Asterisks and brackets indicate significance (ns = not significant, * < 0.05, ** < 0.01). Images and measurement representative of at least 15 samples. Scale bars = 50 µm.

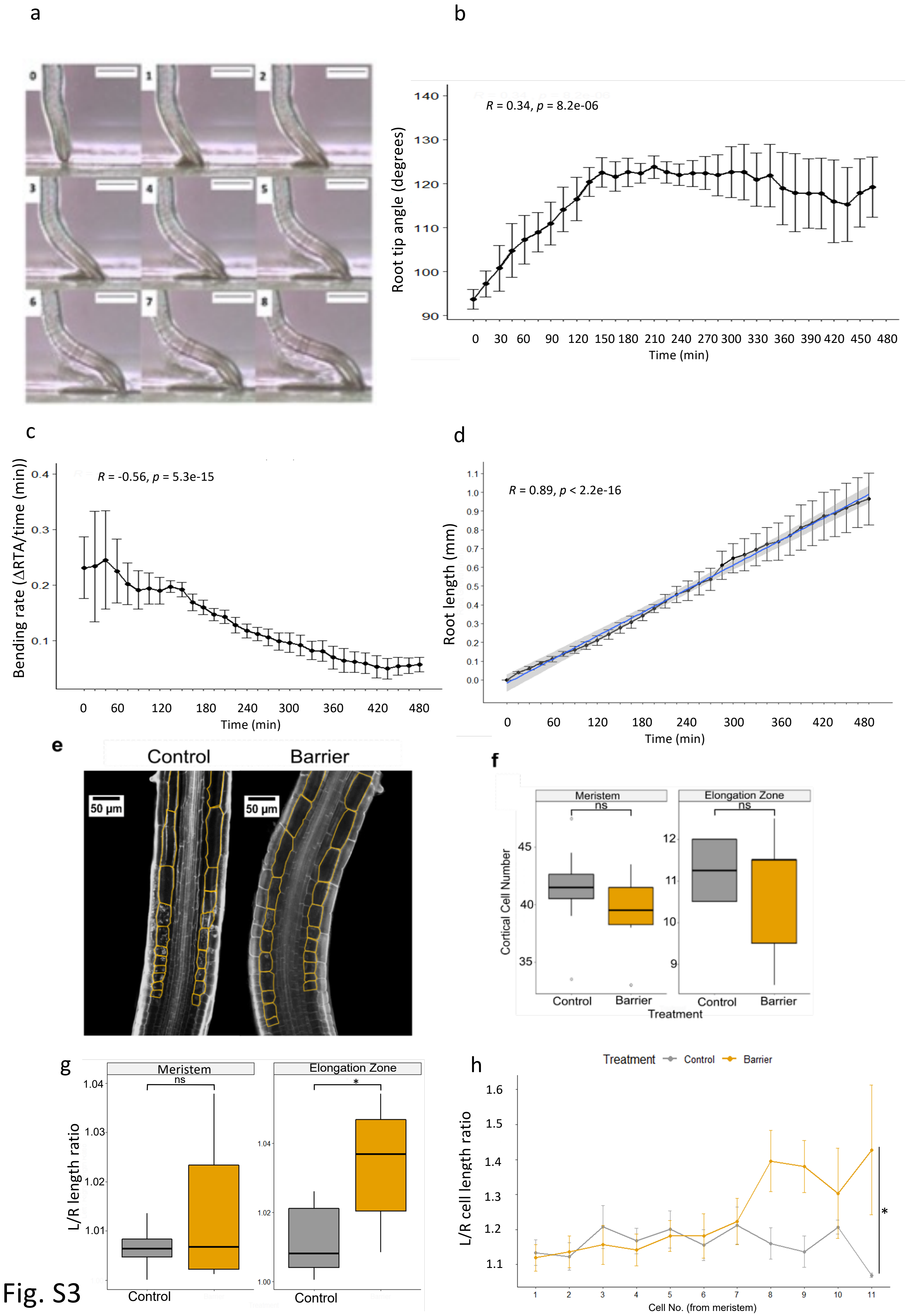
a
b
R = 0.34, p = 8.2e-06
Root tip angle (degrees)
Time (min)
c
R = -0.56, p = 5.3e-15
Bending rate (ΔRTA/time (min))
Time (min)
d
R = 0.89, p < 2.2e-16
Root length (mm)
Time (min)
e
Control
Barrier
50 µm
f
Meristem
Elongation Zone
ns
Cortical Cell Number
Control
Barrier
Treatment
g
L/R length ratio
*
h
Treatment
L/R cell length ratio
Cell No. (from meristem)

Fig. S3

**Figure S3. Short-term bending response of a root to a barrier.** Plastic barriers were placed in front of 6 DAG growing roots and root tips were imaged every 15 min. a) Time-lapse images of a Col-0 root tip encountering a horizontal barrier at 0-8 h after contact with the barrier. b) Root Tip Angle (RTA) over 0-480 min after contact with a barrier. RTA was measured every 15 minutes. c) Bending rate between 15-480 min. d) Root growth of the primary root tip between 0-480 min after contact with the barrier. Blue line represents Regression line. e) Typical elongation zone of primary root tips stained with Calcofluor White and grown in the presence or absence of a barrier for 6 h. Cells of the elongation zone used for measurements are outlined and highlighted in orange. f) Total cortical cell number in the meristem and elongation zone after 6 h. g) Ratio of meristem and elongation zone length between the left and right side of the root tip. h) Ratio of cell length between the left and right side of the root tip in the first 11 cells of the elongation zone. The root tip was divided through the middle into a left and right side and measurements taken separately for each side. Ratio was calculated using the formula exp(|log(left/right)|) to account for any bias in assigning left/right. Error bars represent mean ± SE. Scale bar = 50 µm.

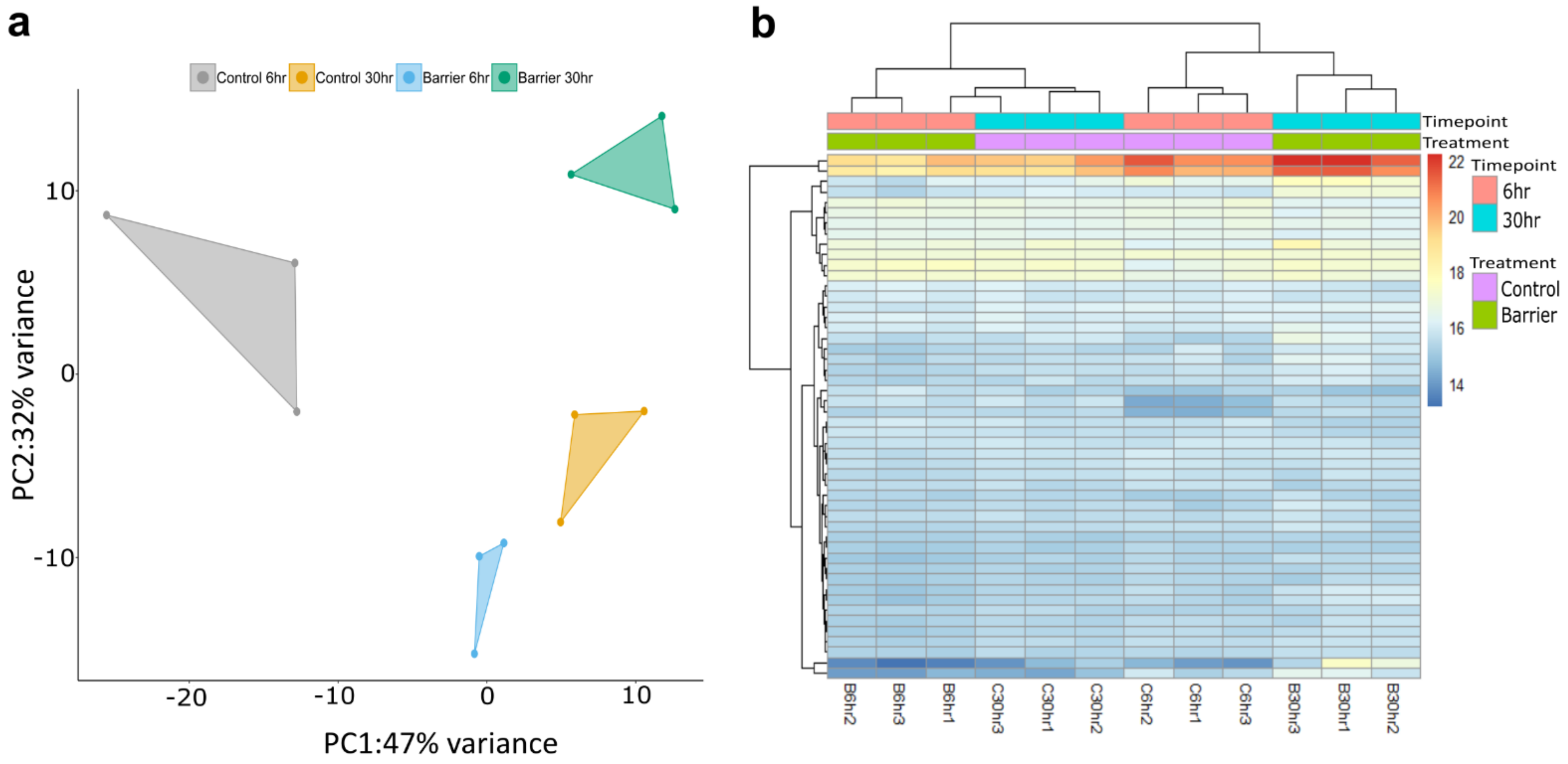


**Fig. S4 Analysis of variation in gene expression between and within sample groups.** a) Principal component analysis (PCA) plot visualising sample-sample distances. Polygons represent sample groups. PCA performed on regularized logarithm (rlog) transformed data using the R software package DEseq2 . b) Clustered Heatmap of rlog transformed count data for the top 50 most highly expressed genes across all samples. Rows and column aggregated using kmeans clustering by the R package pheatmap.

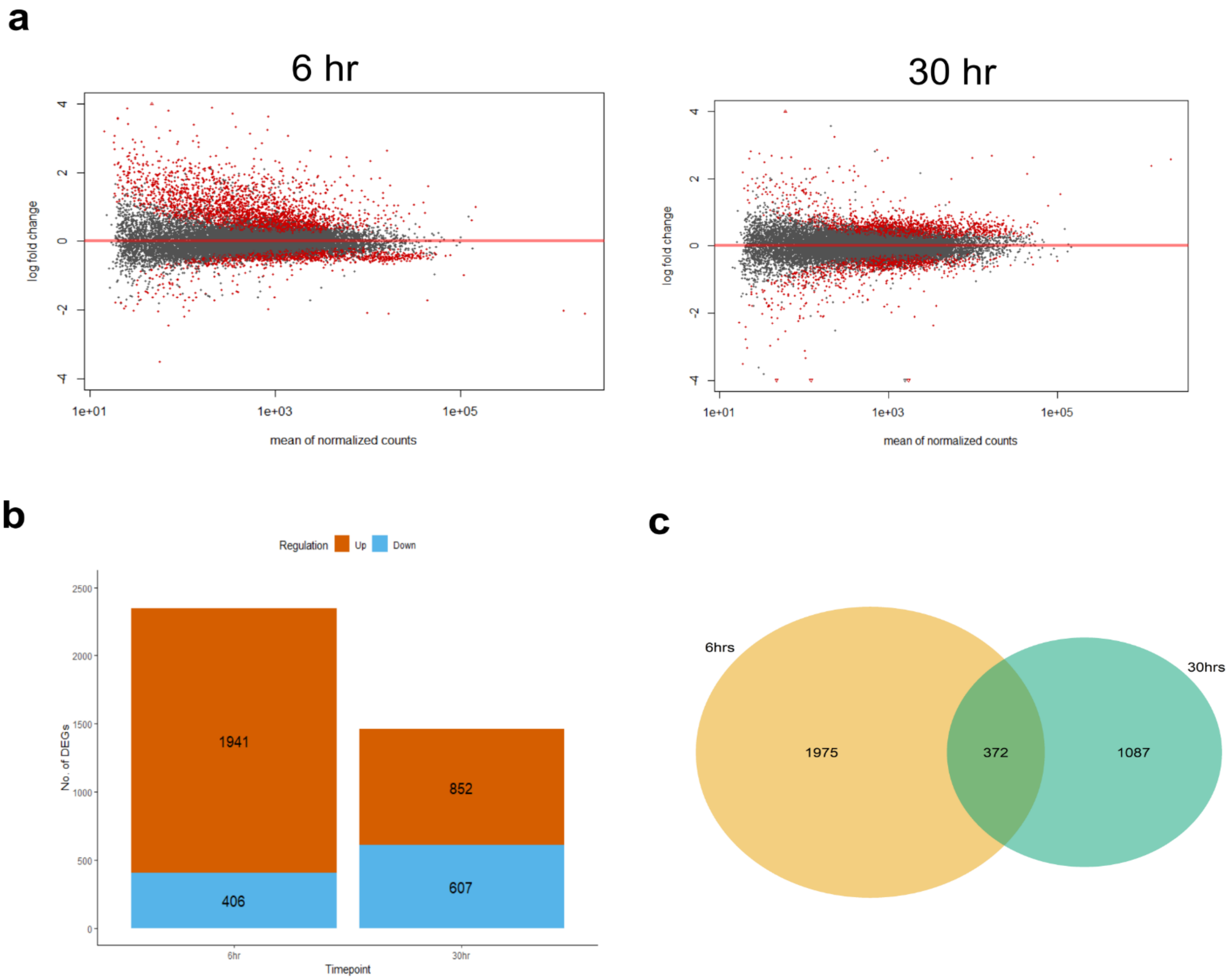
a
6 hr
30 hr
log fold change
mean of normalized counts
1e+01
1e+03
1e+05
b
Regulation
Up
Down
No. of DEGs
1941
406
852
607
6hr
30hr
Timepoint
c
6hrs
30hrs
1975
372
1087

Fig. S5

**Figure S5. Genes differentially expressed in roots encountering a barrier compared with controls after 6 h and 30 h treatment identified through RNA-Seq.**

a) Differentially Expressed Genes (DEGs) as estimated by the R software package DEseq2. Red dots represent significant DEGs, black dots non-significant. P-value < 0.05 and a log2 fold change (log2fc) > 0.5 or < - 0.5. b) Significant DEGs separated into up and downregulated genes at the two barrier response timepoints. c) Venn diagram representing overlapping DEGs between treatments.

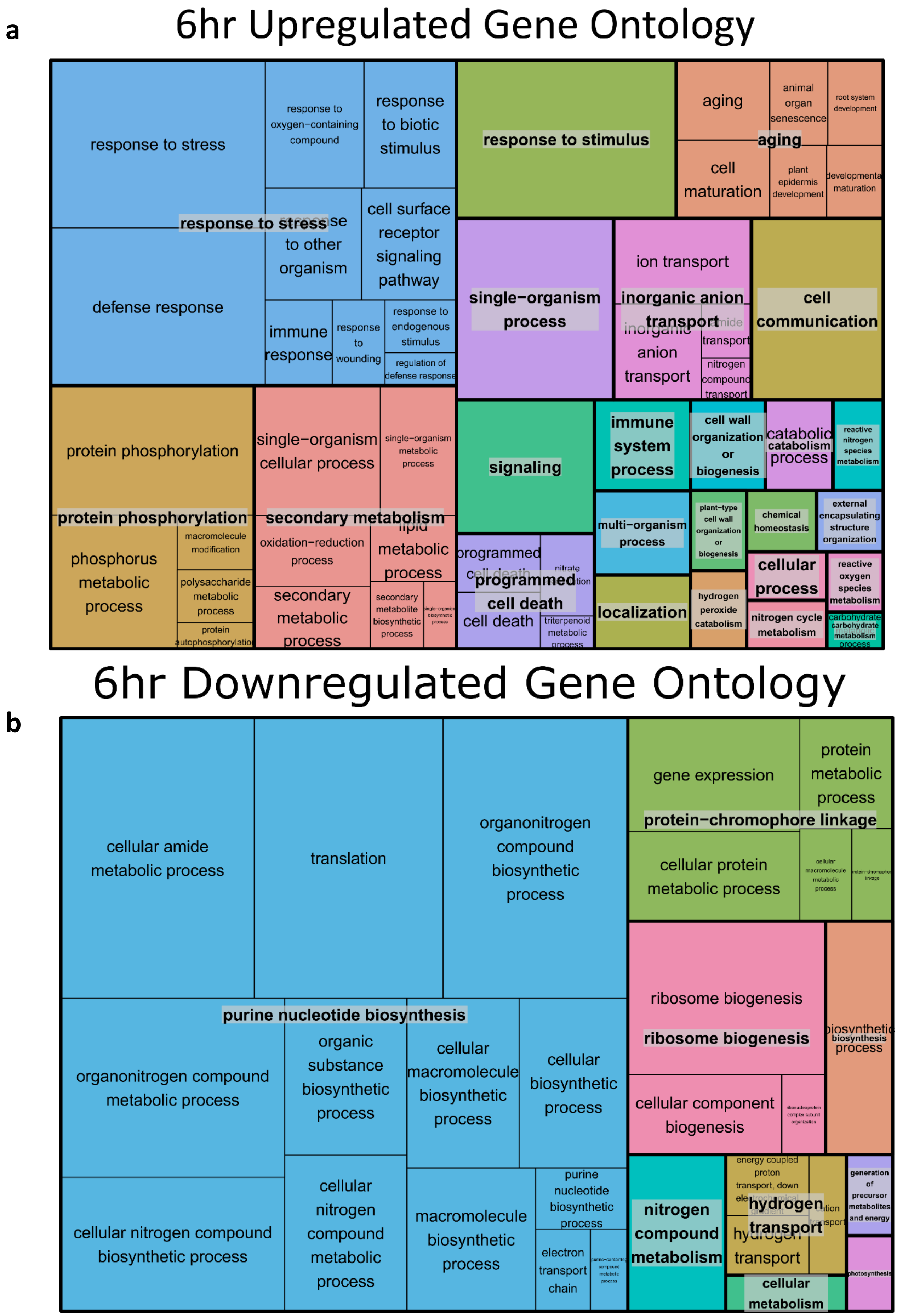


**Figure S6.** Treemap output from REVIGO (Supek et al., 2011) of genes identified as significantly upregulated **(a)** and downregulated **(b)** 6 hours after encountering a barrier. P-value < 0.05 and a log2 fold change (log2fc) > 0.5 or < -0.5. Each rectangle represents a gene ontology (GO) term cluster and each colour represents a supercluster of related clusters. Sizes of rectangles reflect the –log10 P-value of each cluster.

a

### 30hr Upregulated Gene Ontology

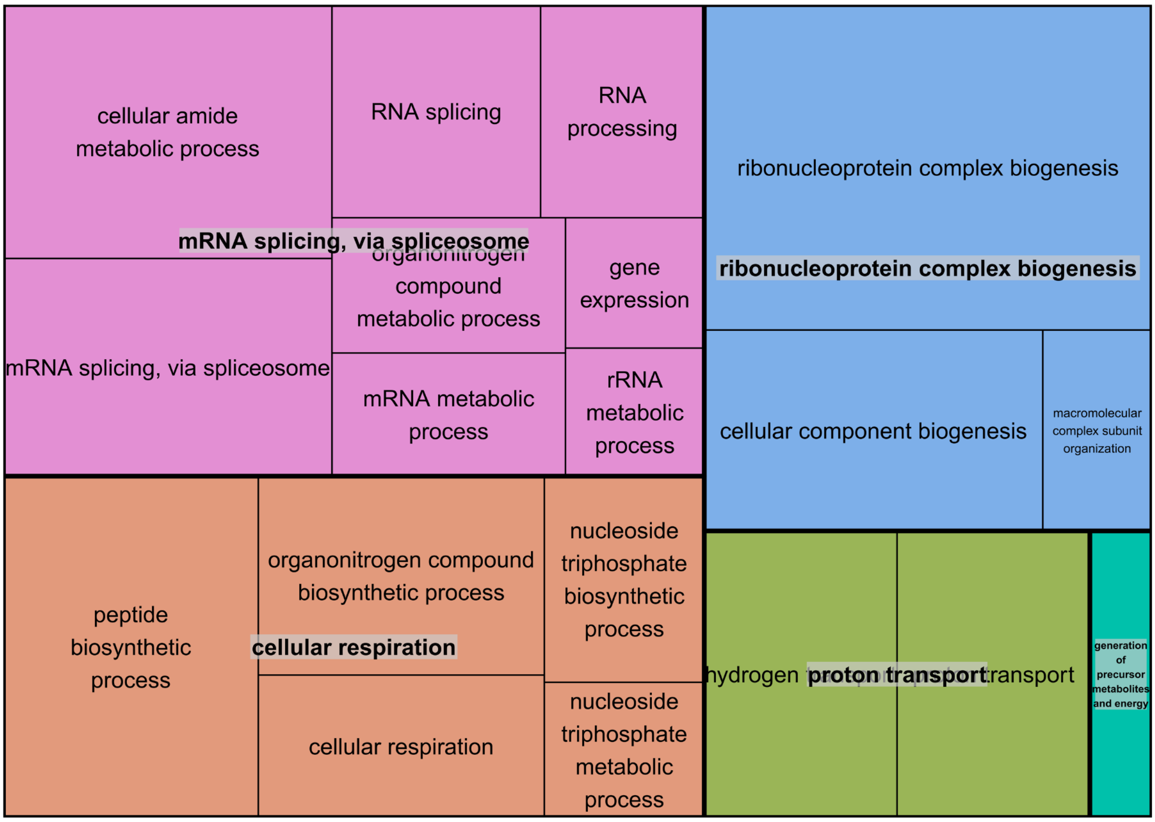


b

### 30hr Downregulated Gene Ontology

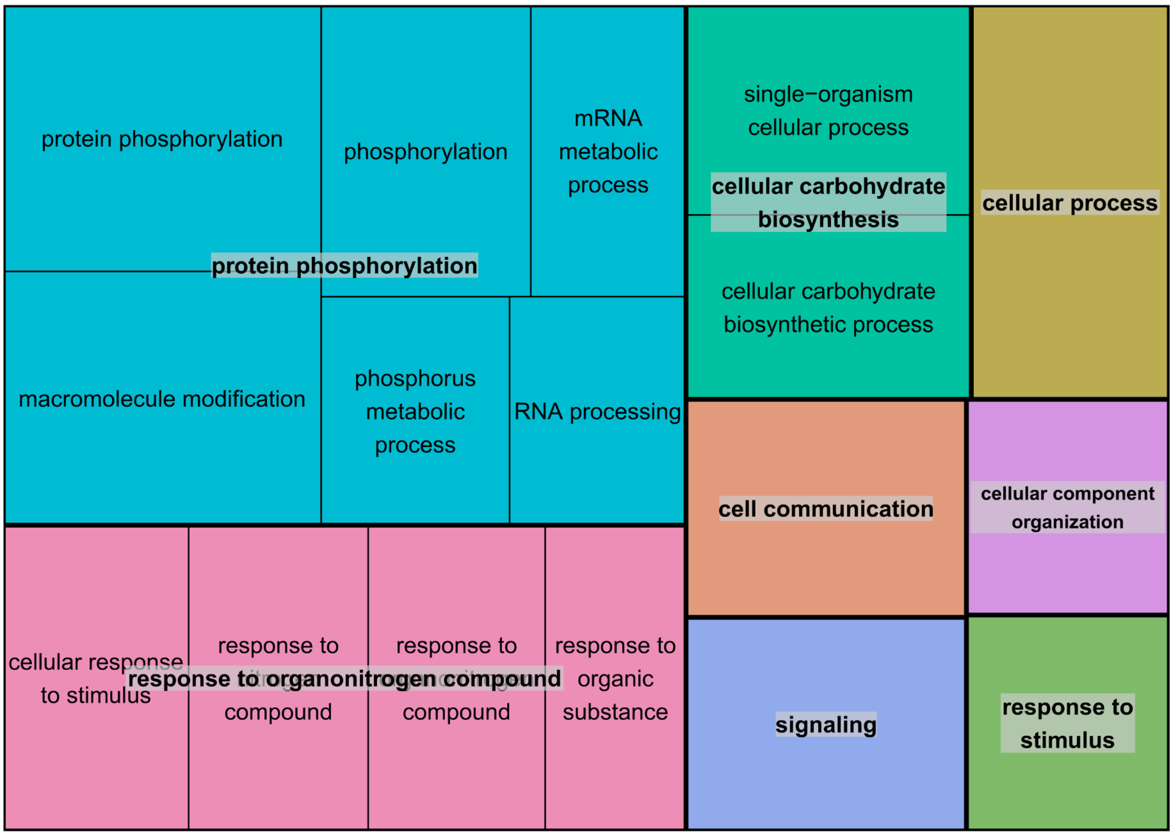


**Figure S7.** Treemap output from REVIGO (Supek et al., 2011) of genes identified as significantly upregulated **(a)** and downregulated **(b)** 30 hours after encountering a barrier. P-value < 0.05 and a log2 fold change (log2fc) > 0.5 or < -0.5. Each rectangle represents a gene ontology (GO) term cluster and each colour represents a supercluster of related clusters. Sizes of rectangles reflect the –log10 P-value of each cluster.

a

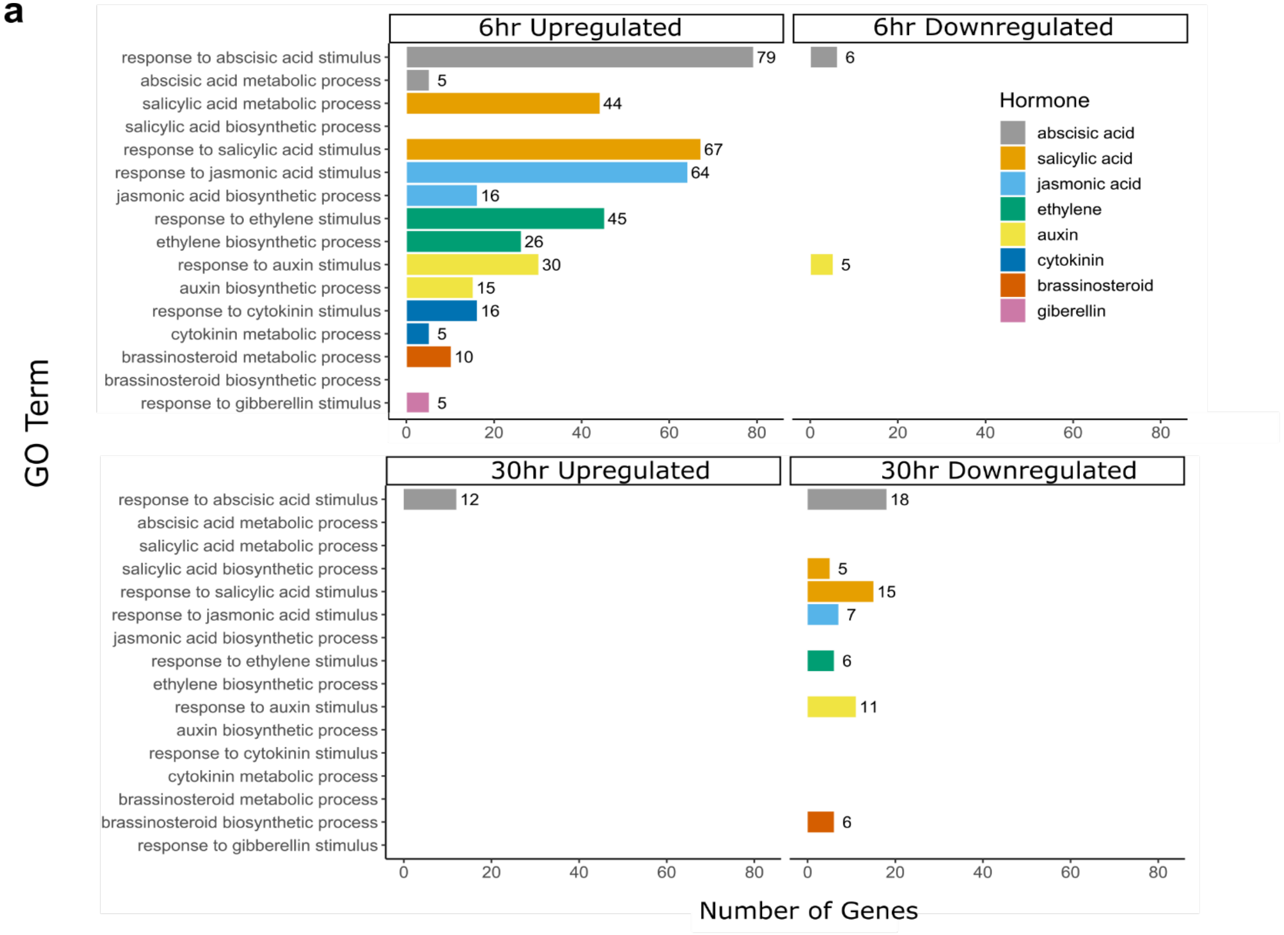


**Figure S8. Hormone signalling and metabolic/biosynthesis related GO terms identified by GO analysis of genes differentially expressed in response to a barrier**. Bar chart showing numbers of DEGs identified by each GO term. Bars are grouped by the hormone the GO term relates to.

**a**

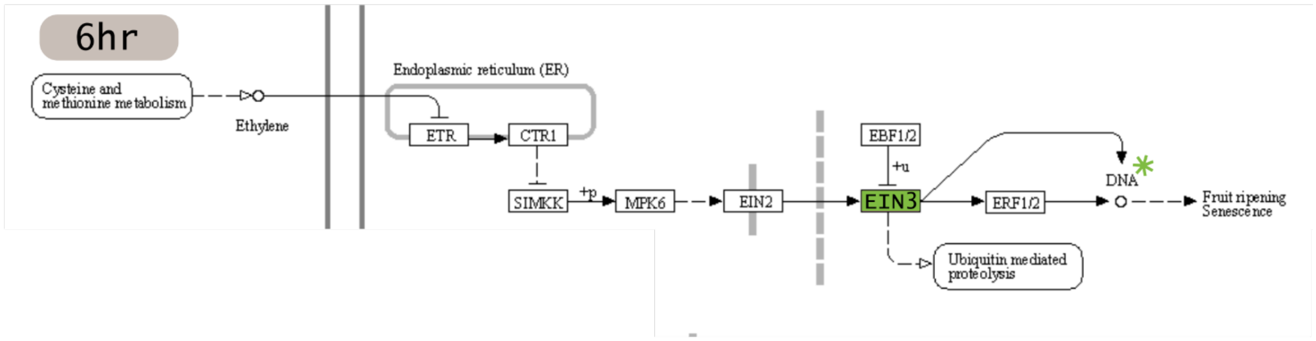


| Locus ID | Gene Symbol | LogFC | KEGG Orthology ID |
|---|---|---|---|
| AT5G21120 | EIL2 | 2.092 | ethylene-insensitive protein 3 (EIN3) |

| Locus ID | Gene Symbol | LogFC | Gene description |
|---|---|---|---|
| AT4G17500 | ERF-1 * | 0.959 | Encodes a member of the ERF (ethylene response factor) subfamily B-3 of ERF/AP2 transcription factor family (ATERF-1). |
| AT3G15210 | ERF4 * | 0.769 | Encodes a member of the ERF (ethylene response factor) subfamily B-1 of ERF/AP2 transcription factor family (ATERF-4). |

**b**

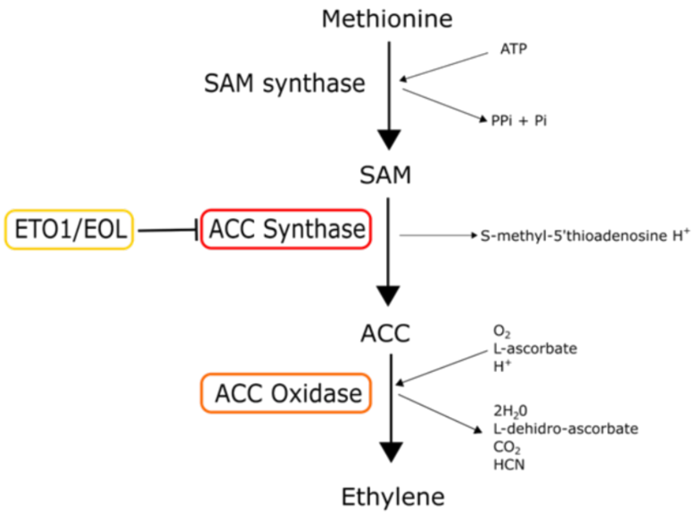


| Locus ID | Gene Symbol | LogFC 6hr | LogFc 30hr | Gene Description |
|---|---|---|---|---|
| AT5G65800 | ACC SYNTHASE 5 (ACS5) | 2.520 | -1.02 | 1-aminocyclopropane-1-carboxylate synthase (ACS) is encoded by a multigene family consisting of at least five members whose expression is induced by hormones, developmental signals, and protein synthesis inhibition |
| AT1G05010 | ETHYLENE-FORMING ENZYME (EFE) | 0.794 | 0 | Encodes 1-aminocyclopropane-1-carboxylate oxidase |
| AT5G58550 | ETO1-LIKE 2 (EOL2) | 0.697 | -1.5 | Encodes a paralog of ETO1, which is a negative regulator of ACS5 (a key enzyme in ethylene biosynthesis pathway). EOL2 also interacts with and inhibits the activity of ACS5 |

**Figure S9. Ethylene-related gene expression analysis.** a) KEGG Pathway mapping of genes differentially expressed at 6 hours in response to a barrier and identified as being involved in the ethylene signalling pathway. Genes present within the data set are highlighted with colours corresponding to KEGG Orthology (molecular function) definition. b) Genes differentially expressed at 6 hours in response to a barrier and identified as being involved in the ethylene biosynthesis pathway. Components of the pathway with an identified DEG are highlighted red (ACC Synthase), orange (ACC Oxidase) and yellow (ETO1/EOL1).

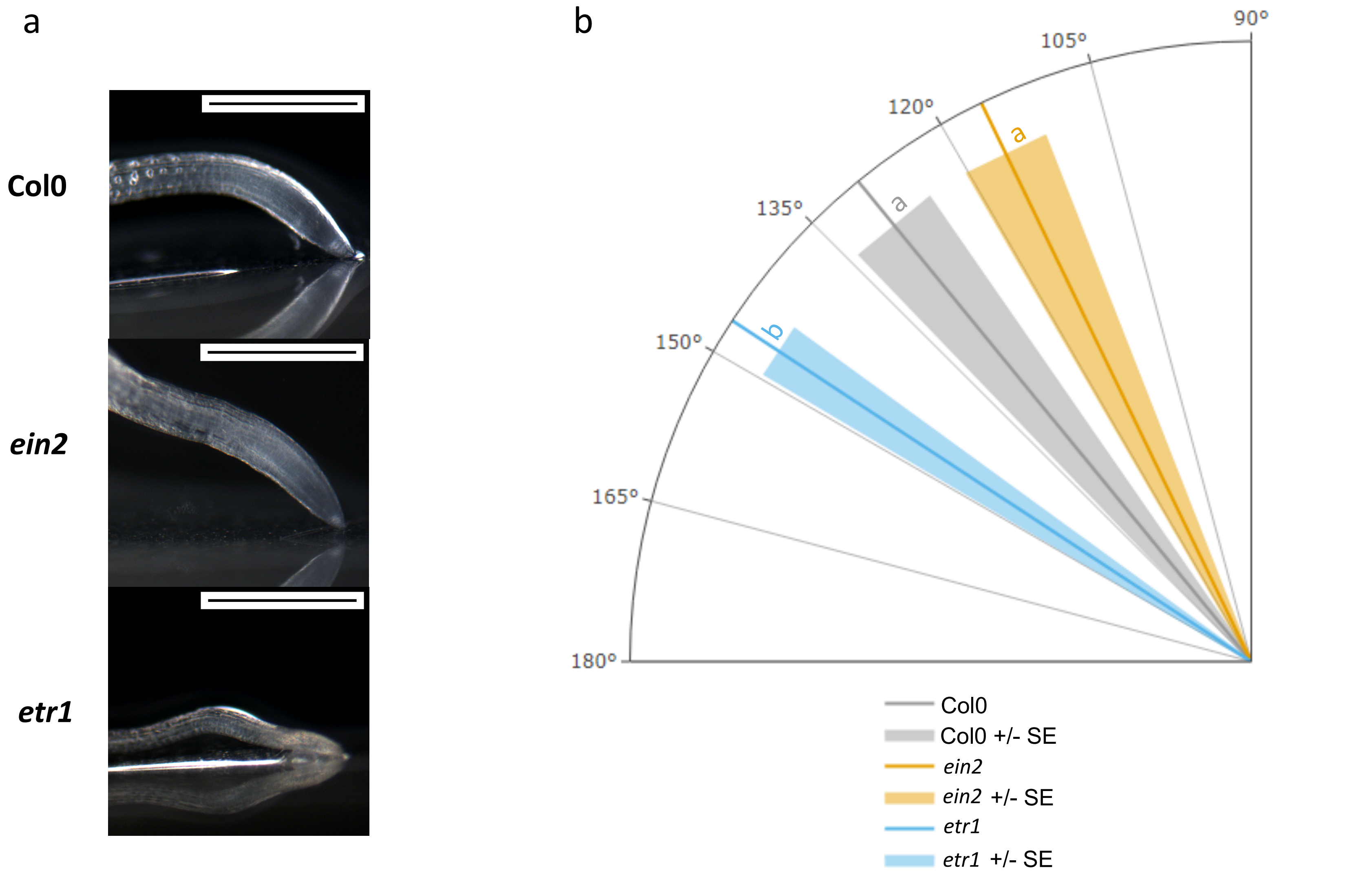


**Figure S10. Response of *etr1* and *ein2* to a barrier.** a) Plastic barriers were placed in front of primary roots of seedlings 6 days after germination (DAG) and root tips were imaged at 24 hours after encountering a barrier. Scale bar indicates 0.5 mm. b) Angle of primary root tips to the horizontal barrier. Lines indicate mean and surrounding shaded area indicates ± SE. Letters indicate significance with a Tukey Pairwise comparison $P < 0.05$

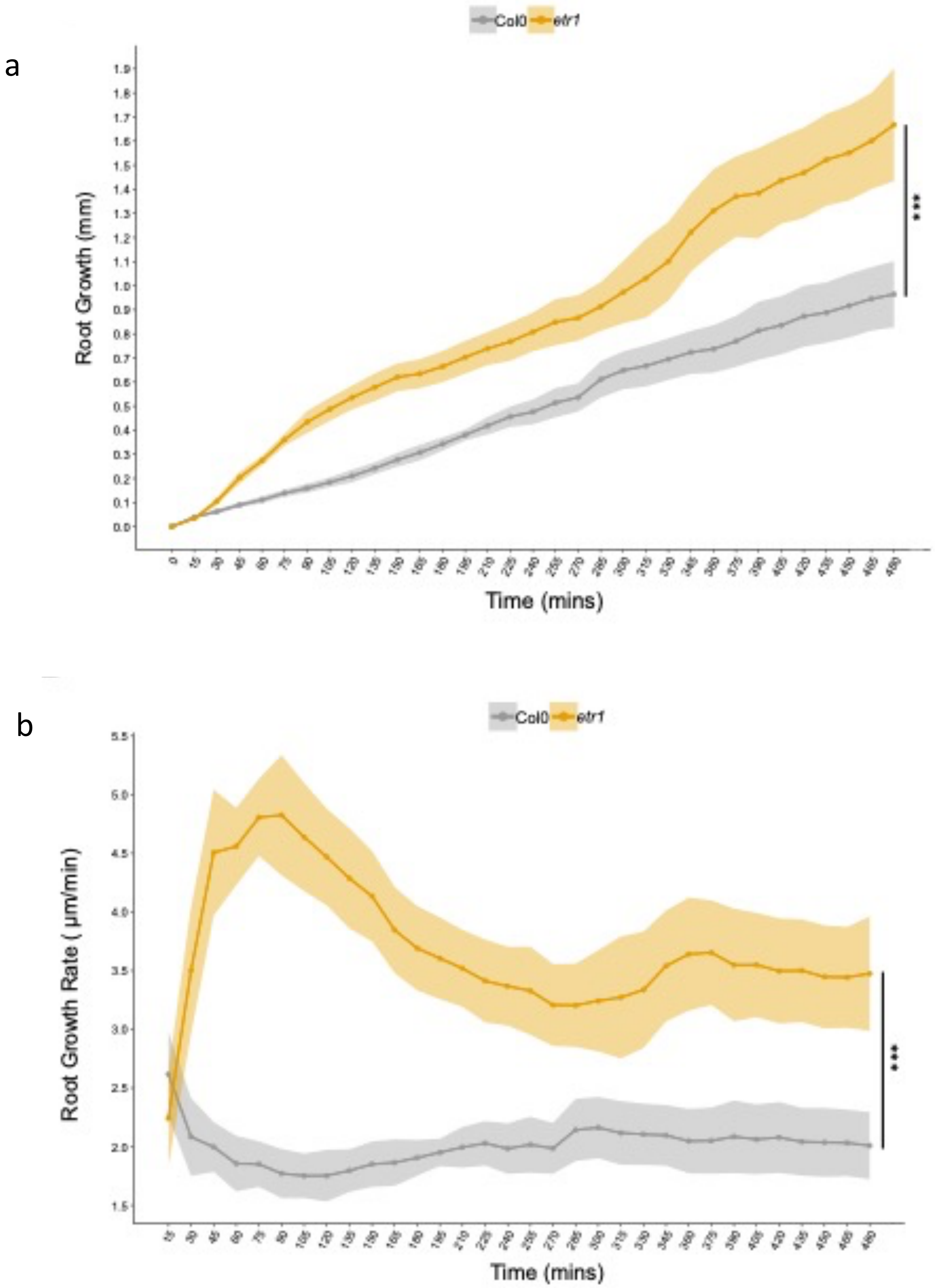


**Figure S11. Growth of *etr1* between 0 and 8 hours after barrier placement.** Root growth as measured by time-lapse imaging of roots encountering a barrier. a) Root growth of the primary root tip between 0-480 minutes after contact with the barrier. b) Root growth rate (µm/min) between 15-480 minutes after encountering a barrier. Plastic barriers were placed in front of 6 day old vertically growing roots and root tips were imaged every 15 min. Lines and dots indicate mean with shaded area indicating ± SE. Letters indicate significance after a *Student's* t-test ( *** < 0.001).

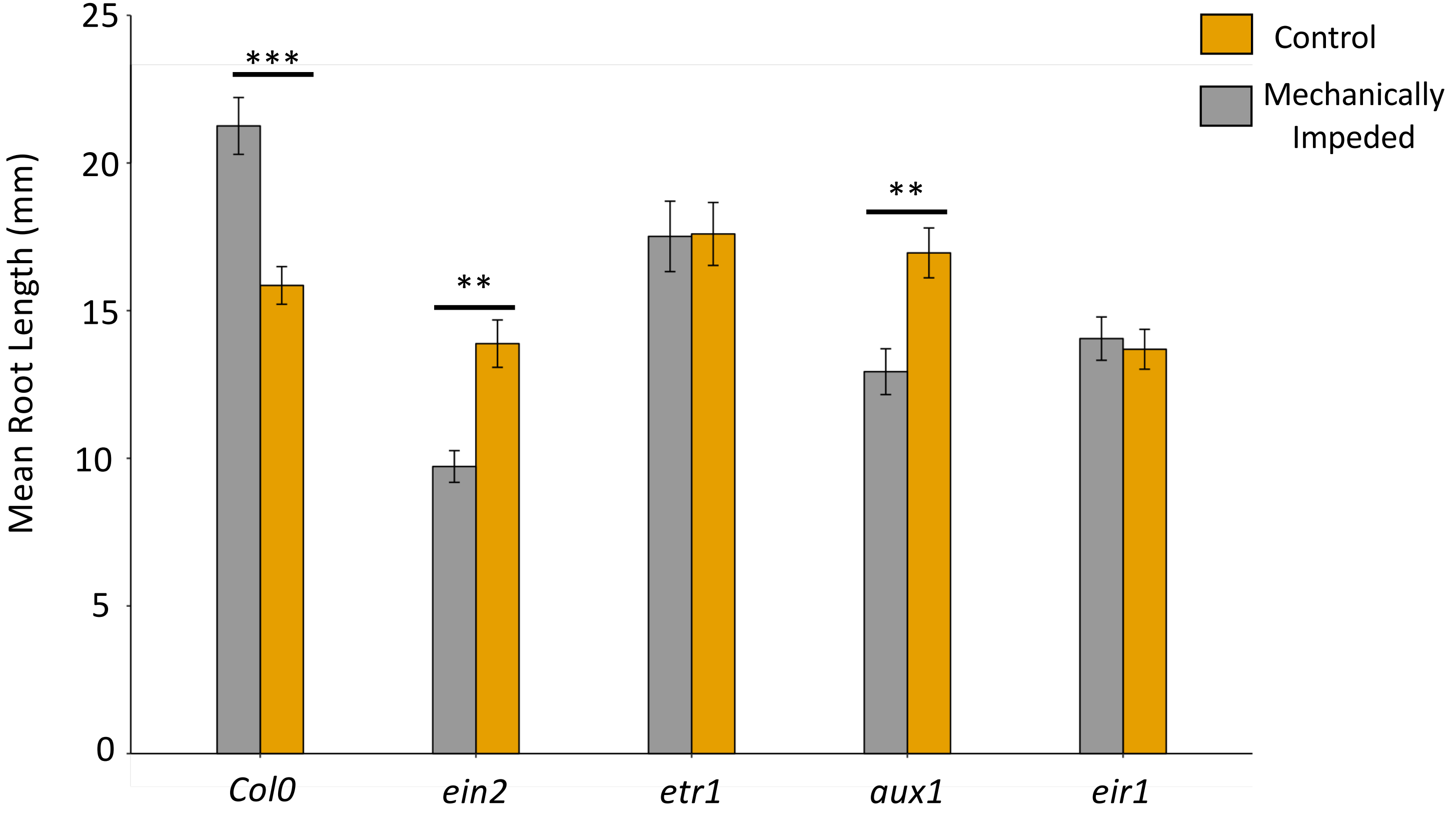


**Figure S12. Growth of wildtype and ethylene-sensitive mutant roots after barrier placement.** Primary root length was measured after growth of seedlings on dialysis membrane barriers for 7 days post germination. Error bars represent standard error of the mean of 15 seedlings per treatment. Root growth of the mutants was not significantly inhibited by barrier impedance (ANOVA and Tukey pairwise comparison, *** = P <0.0001, ** + P <0.001).

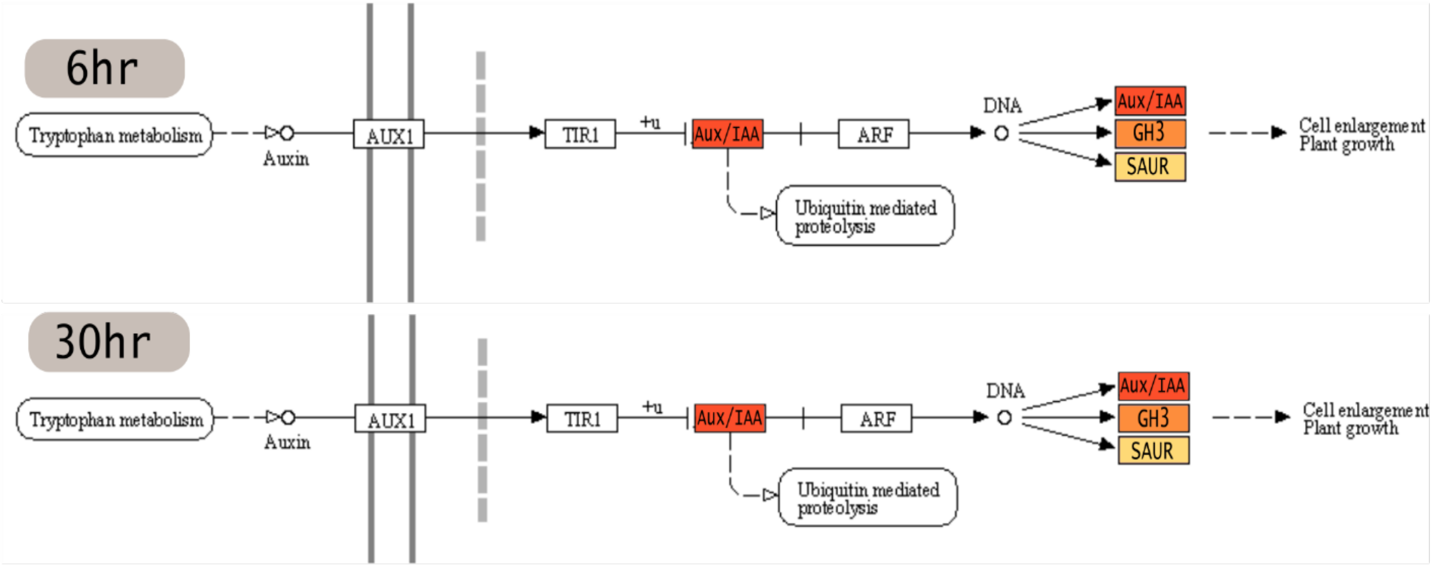


6hr

| Locus ID | Gene Symbol | LogFC | KEGG Orthology ID |
|---|---|---|---|
| AT4G29080 | PAP2 | -0.58 | auxin-responsive protein IAA **(AUX/IAA)** |
| AT1G2830 | GH3.17 | 1.18 | auxin-responsive GH3 gene family **(GH3)** |
| AT4G37390 | BRU6 | 1.19 | auxin-responsive GH3 gene family **(GH3)** |
| AT5G54510 | DFL1 | 0.65 | auxin-responsive GH3 gene family **(GH3)** |
| AT2G45210 | SAUR36 | -0.77 | SAUR family protein **(SAUR)** |
| AT5G50760 | SAUR55 | 0.65 | SAUR family protein **(SAUR)** |

30hr

| Locus ID | Gene Symbol | LogFC | KEGG Orthology ID |
|---|---|---|---|
| AT4G29080 | IAA30 | -0.58 | auxin-responsive protein IAA **(AUX/IAA)** |
| AT1G2830 | IAA14 | 1.18 | auxin-responsive protein IAA **(AUX/IAA)** |
| AT4G37390 | PAP2 | 1.19 | auxin-responsive protein IAA **(AUX/IAA)** |
| AT5G54510 | GH3.17 | 0.65 | auxin-responsive GH3 gene family **(GH3)** |
| AT2G45210 | DFL2 | -0.77 | auxin-responsive GH3 gene family **(GH3)** |
| AT5G50760 | SAUR36 | 0.65 | SAUR family protein **(SAUR)** |

**Figure S13. KEGG Pathway mapping of genes differentially expressed at 6 and 30 hours in response to a barrier and identified as being involved in the auxin signalling pathway.** Genes present within the data set are highlighted with colours corresponding to KEGG Orthology (molecular function) definition.

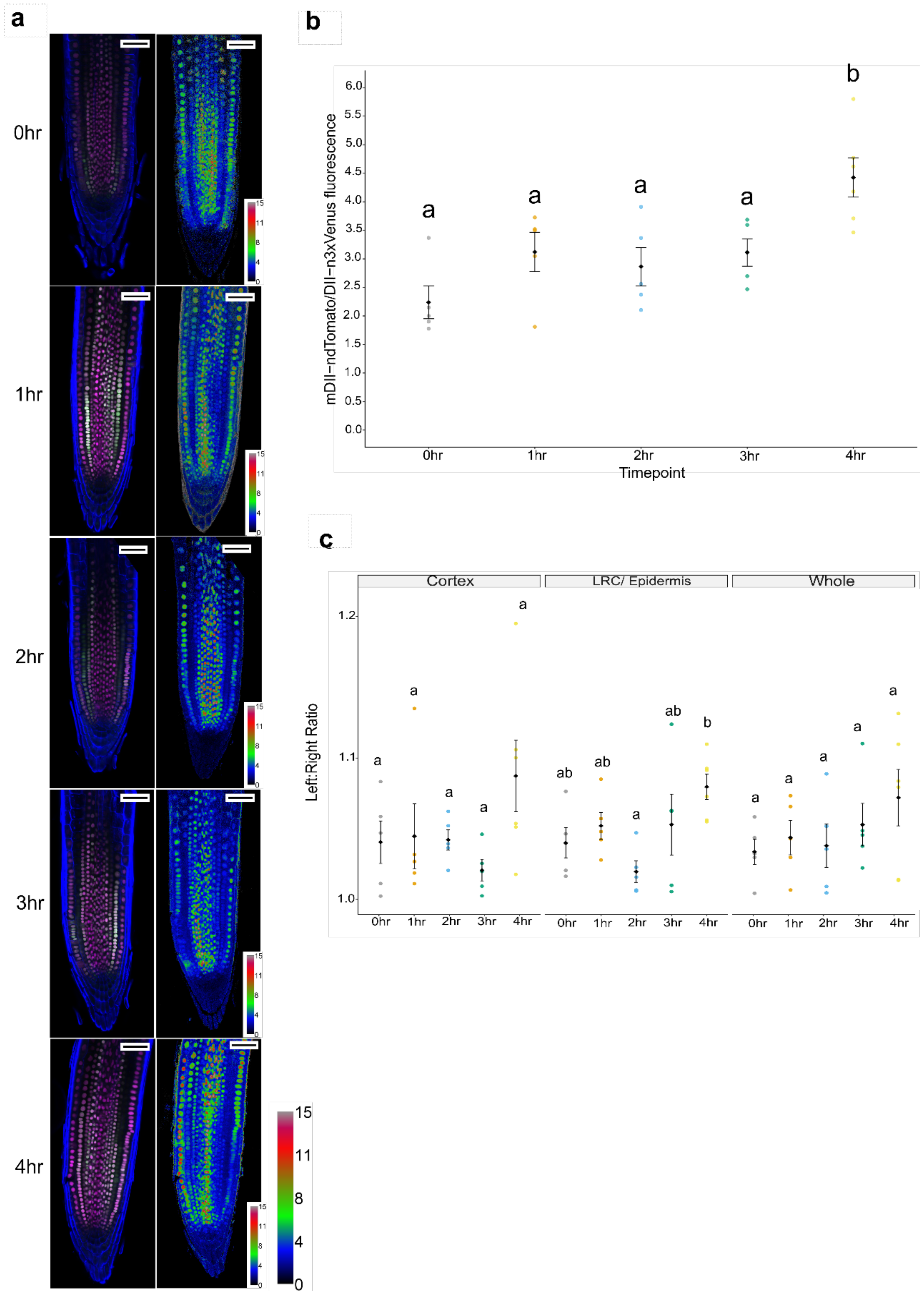


**Figure S14. Confocal imaging of R2D2 in roots responding to a barrier between 0-4 h**. Barriers were placed in front of root tips 6 DAG. a) R2D2 fluorescence at the root tip at 0-4 hours with accompanying ratio metric image of generated from mDII-ndTomato/ DII-m3xVenus fluorescence using ImageJ. For ratiometric images calibration bar indicates mean grey value. b) Ratio of mDII-ndTomato/ DII-m3xVenus fluorescence. c) Ratio of auxin level across the left and right sides of the root tip. Scale bar = 50 µm. Black circles and error bars represent mean ± SE. Coloured circles represent distribution of individual data points. Letters indicate significant difference after post-hoc TUKEY test, $P < 0.05$.

**Supplemental Table 1** NADPH-oxidase genes identified through RNA-Seq that are upregulated during the barrier response at 6 hours.

| Locus ID | Primary Gene Symbol | Log FC 6hr | Gene Description |
|---|---|---|---|
| AT1G09090 | RBOHB | 0.608 | NADPH-oxidase AtrbohB plays a role in seed after-ripening. Major producer of superoxide in germinating seeds. AtrbohB pre-mRNA is alternatively spliced in seeds in a hormonally and developmentally regulated manner. |
| AT1G64060 | RBOHF | 0.608 | Interacts with AtrbohD gene to fine tune the spatial control of ROI production and hypersensitive response to cell in and around infection site. |
| AT4G11230 | RBOHI | 0.694 | NADPH-oxidase RbohI is expressed highly in seeds and roots. Mutants have inreased sensitivity to osmotic stress suggesting a role in mediating cellular response to stress in roots. |
| AT5G07390 | RBOHA | 1.063 | respiratory burst oxidase homolog A |
| AT5G47910 | RBOHD | 0.743 | NADPH/respiratory burst oxidase protein D (RbohD).Interacts with AtrbohF gene to fine tune the spatial control of ROI production and hypersensitive response to cell in and around infection site. |
| AT5G51060 | RHD2 | 1.145 | RHD2 (along with RHD3 and RHD4) is required for normal root hair elongation. Has NADPH oxidase activity. Gene is expressed in the elongation and differention zone in trichoblasts and elongating root hairs. Required for the production of reactive oxygen species in response to extracellular ATP stimulus. The increase in ROS production stimulates $Ca^{2+}$ influx. |

**Supplemental Table 2** List of genes that act as reactive oxygen species (ROS) scavengers identified in the RNA-Seq data of differentially expressed genes in response to a barrier log2fc identified with P-value <0.05.

| Locus ID | Gene Symbol | Enzyme Family | LogFC |
|---|---|---|---|
| AT1G01580 | FERRIC REDUCTION OXIDASE 2 (FRO2) | NADPH oxidase-like | 1.918 |
| AT1G03850 | GLUTAREDOXIN 13 (GRXS13) | Glutaredoxin (GLR) | 0.970 |
| AT1G09090 | RESPIRATORY BURST OXIDASE HOMOLOG B (RBOHB) | NADHP oxidase | 0.608 |
| AT1G19230 | (ATRBOHE) | NADPH oxidase | 0.977 |
| AT1G23020 | FERRIC REDUCTION OXIDASE 3 (FRO3) | NADPH oxidase | 0.849 |
| AT1G28480 | (GRX480) | Glutaredoxin (GLR) | 1.931 |
| AT1G45145 | THIOREDOXIN H-TYPE 5 (TRX5) | Thioredoxins (Trx) | 0.760 |
| AT1G60740 | | Peroxiredoxin (PrxR) | 0.883 |
| AT1G64060 | RESPIRATORY BURST OXIDASE PROTEIN F (RBOH F) | NADPH oxidase | 0.608 |
| AT2G31570 | GLUTATHIONE PEROXIDASE 2 (GPX2) | Glutathione Peroxidase (GPX) | 0.804 |
| AT3G27820 | MONODEHYDROASCORBATE REDUCTASE 4 (MDAR4) | Monodehydroascorbate Reductase (MDAR) | 0.523 |
| AT4G11230 | (RBOHI) | NADPH oxidase | 0.694 |
| AT4G25090 | (ATRBOHG) | NADPH oxidase | 1.595 |
| AT4G35970 | ASCORBATE PEROXIDASE 5 (APX5) | Ascorbate Peroxidase (APX) | 0.855 |
| AT5G07390 | RESPIRATORY BURST OXIDASE HOMOLOG A (RBOHA) | NADPH oxidase | 1.063 |
| AT5G23980 | FERRIC REDUCTION OXIDASE 4 (FRO4) | NADPH oxidase-like | 1.680 |
| AT5G23990 | FERRIC REDUCTION OXIDASE 5 (FRO5) | NADPH oxidase-like | 2.451 |
| AT5G36270 | | Dehydroascorbate Reductase (DHAR) | 2.578 |
| AT5G47910 | RESPIRATORY BURST OXIDASE HOMOLOGUE D (RBOHD) | NADPH oxidase | 0.743 |
| AT5G51060 | ROOT HAIR DEFECTIVE (RHD2) | NADPH oxidase | 1.145 |
| AT3G62930 | GRXS6 | Glutaredoxin (GLR) | -0.658 |
